# LJA: Assembling Long and Accurate Reads Using Multiplex de Bruijn Graphs

**DOI:** 10.1101/2020.12.10.420448

**Authors:** Anton Bankevich, Andrey Bzikadze, Mikhail Kolmogorov, Dmitry Antipov, Pavel A. Pevzner

## Abstract

Although most existing genome assemblers are based on the de Bruijn graphs, it remains unclear how to construct these graphs for large genomes and large *k*-mer sizes. This algorithmic challenge has become particularly important with the emergence of long high-fidelity (HiFi) reads that were recently utilized to generate a semi-manual telomere-to-telomere assembly of the human genome and to get a glimpse into biomedically important regions that evaded all previous attempts to sequence them. To enable automated assemblies of long and accurate reads, we developed a fast LJA algorithm that reduces the error rate in these reads by three orders of magnitude (making them nearly error-free) and constructs the de Bruijn graph for large genomes and large *k*-mer sizes. Since the de Bruijn graph constructed for a fixed *k*-mer size is typically either too tangled or too fragmented, LJA uses a new concept of a multiplex de Bruijn graph with varying *k*-mer sizes. We demonstrate that LJA improves on the state-of-the-art assemblers with respect to both accuracy and contiguity and enables automated telomere-to-telomere assemblies of entire human chromosomes.

## Introduction

The emergence of long and accurate reads opened a possibility to generate the first complete (telomere-to-telomere) assembly of a human genome and to get a glimpse into biomedically important genomic regions that evaded all previous attempts to sequence them (Nurk et al., 2021). The Telomere-to-Telomere (T2T) and the Human Pangenome Reference (HPR) Projects are now using long and accurate reads for population-scale assembly of multiple human genomes and for diagnosing rare diseases that remained below the radar of short-read technologies (Ebert et al., 2021, Hiatt et al., 2021, Miller et al., 2021).

These breakthroughs in genome sequencing were mainly achieved using *high-fidelity* (*HiFi*) reads (Wenger et al., 2019). However, assembly of HiFi reads is far from being straightforward: the complete assembly of a human genome was generated using a semi-manual effort of a large consortium rather than by an automated approach (Nurk et al., 2021). Since such time-consuming efforts are neither sustainable nor scalable in the era of population-scale sequencing, there is a need for an accurate (nearly error-free) tool for complete genome assembly.

We argue that this challenge requires an algorithm for constructing *large de Bruijn graphs* (Compeau et al., 2011), i.e., de Bruijn graphs for large genomes and large *k*-mer sizes exceeding a thousand of nucleotides. Indeed, similarly to assembling short and accurate reads, the de Bruijn graph approach has the potential to improve assemblies of any type of accurate reads. However, although the de Bruijn graphs represent the algorithmic engine of nearly all short-read assemblers (Zerbino and Birney, 2008, Simpson et al., 2009, Peng et al., 2010, Bankevich et al., 2012), the problem of constructing large de Bruijn graphs remains open and the existing HiFi assemblers HiCanu (Nurk et al., 2020) and hifiasm (Cheng et al., 2021) are based on the alternative *string graph* approach (Myers, 2005).

Since HiFi reads are on average even more accurate than Illumina reads (Wenger et al., 2019), the de Bruijn graph approach is expected to work well for their assembly (see Lin et al. 2014 for a comparison of the de Bruijn graphs and the string graphs). Application of this approach to long HiFi reads requires either constructing the de Bruijn graph with a large *k*-mer size or alternatively, using the de Bruijn graph with a small *k*-mer-size for follow-up repeat resolution by threading long reads through this graph (Antipov et al., 2016). However, it remains unclear how to address three open algorithmic problem in assembling long and accurate reads: (i) constructing large de Bruijn graphs, (ii) error-correcting HiFi reads so that they become nearly error-free and thus amenable to applying the de Bruijn graph approach, and (iii) utilizing the entire length of long reads for resolving repeats that are longer than the *k*-mer size.

The existing genome assemblers are not designed for constructing large de Bruijn graphs since their time/memory requirements become prohibitive when the *k*-mer size becomes large, e.g., simply storing all 5001-mers of the human genome requires ≅4 TB of memory. For example, the SPAdes assembler (Bankevich et al., 2012) faces time/memory bottlenecks assembling mammalian genomes with the *k*-mer size exceeding 500. To reduce the memory footprint, some assembly algorithms avoid explicitly storing all *k*-mers by constructing a *perfect hash map* (Fredman et al., 1984, Bankevich et al., 2012) or the *Burrows-Wheeler Transform* (Burrows and Wheeler, 1984, Li et al., 2015) of all reads. However, even with these improvements, the memory footprint remains prohibitively large, not to mention that the running time is proportional to the *k*-mer size.

The *repeat graph* approach (Pevzner et al., 2004) and the *sparse de Bruijn graph* approach (Ye et al., 2012) construct coarse versions of the de Bruijn graph with smaller time/memory requirements. Recently, Kolmogorov et al., 2020 modified the Flye assembler for constructing the repeat graph of HiFi reads (in a metagenomic context), while Rautiainen and Marschall, 2021 showed how to assemble HiFi reads into a sparse de Bruijn graph. However, these graphs represent coarse versions of the de Bruijn graph, thus limiting their capabilities in assembling the highly-repetitive regions such as centromeres.

We introduce *La Jolla Assembler* (*LJA*) that includes three modules addressing all three challenges in assembling long and accurate reads: jumboDBG (constructing large de Bruijn graphs), mowerDBG (error-correcting reads), and multiplexDBG (utilizing the entire read-length for resolving repeats). jumboDBG combines four algorithmic ideas: the *Bloom filter* (Bloom, 1970), the *rolling hash* (Karp and Rabin, 1987), the sparse de Bruijn graph (Ye et al., 2012), and the *disjointig* generation (Kolmogorov et al., 2019). Although each of these ideas was used in previous bioinformatics studies, jumboDBG is the first approach that combines them. LJA launches jumboDBG to construct the de Bruijn graph, launches mowerDBG that uses this graph to correct nearly all errors in reads, launches jumboDBG again to generate a much simpler graph of the error-corrected reads, and launches multiplexDBG to transform it into the *multiplex de Bruijn graph* with varying *k*-mer sizes to take advantage of the entire read-lengths. LJA also includes the LJApolish module that expands the collapsed homopolymer runs in the resulting assembly.

Although we benchmarked LJA, hifiasm, and HiCanu on various mammalian, plant, and insect genomes, evaluating the quality of the resulting assemblies is challenging since neither the complete reference for these genomes nor an automated pipeline for a reference-grade assembly validation are available yet (Mc Cartney et al., 2021). We thus focused on benchmarking these assemblers using the HiFi read-set (referred to as the T2T dataset) from a haploid human CHM13 cell line assembled by the T2T consortium (Nurk et al., 2021). This painstakingly validated assembly represents the only accurate telomere-to-telomere sequence of a large genome available today. LJA generated the most contiguous assembly of this dataset, (including complete assemblies of six human chromosomes without any misassemblies and only ten misassemblies for the entire human genome), reducing the number of assembly errors five-fold as compared to hifiasm and HiCanu. The accuracy of genome assemblers becomes critical in the era of population-wide complete genome sequencing since semi-manual validation of complete genome assemblies (Mc Cartney et al., 2021), is prohibitively time-consuming.

## Results

### Brief description of the algorithmic concepts used in the LJA pipeline

The goal of genome assembly is to reconstruct a genome from its error-prone fragments (*reads*). Given a string-set *Reads* and an integer *k*, the (uncompressed) de Bruijn graph *UDB*(*Reads,k*) is a directed graph where each vertex is a *k*-mer from *Reads* and each (*k*+1)-mer *a_1_a_2_ … a_k_ a_k+1_* in reads corresponds to an edge connecting vertices *a_1_a_2_ … a_k_* and *a_2_ … a_k_ a_k+1_*. In particular, the uncompressed de Bruijn graph of a genome *UDB*(*Genome,k*) is defined by considering each chromosome in *Genome* as a single “read”. We refer to an error-free read-set *Reads* that contains all (*k*+1)-mers from a string-set *Genome* as a *k-complete* read-set and note that *UDB*(*Genome,k*)=*UDB*(*Reads,k*) for a *k*-complete read-set.

A vertex with the indegree *N* and the outdegree *M* is referred to as an *N-in-M-out vertex.* A vertex is *non-branching* if it is a 1-in-1-out vertex, and a *junction*, otherwise. We refer to the set of all junctions (*k*-mers) in the graph *UDB*(*Reads,k*) as *Junctions*(*Reads,k*). A path between junctions is *non-branching* if all its intermediate vertices are non-branching. A set of *k*-mers from *Reads* is called a *junction-superset* if it contains all junctions in *UDB*(*Reads*,*k*).

The *compressed de Bruijn graph DB*(*Reads*,*k*) is a compact version of the uncompressed graph *UDB*(*Reads,k*) where each non-branching path is compressed into an appropriately labeled single edge (see Supplementary Note 1). Since LJA utilizes compressed de Bruijn graphs, we refer to them simply as the “de Bruijn graphs”. The *coverage* of an edge in *UDB*(*Reads,k*) is defined as the number of times the label of this edge occurs in *Reads*. The *coverage* of an edge in *DB*(*Reads*, *k*) is defined as the average coverage of all edges in the non-branching path that was compressed into this edge.

### The challenge of constructing a large de Bruijn graph

Since the compressed de Bruijn graph *DB*(*Genome*,*k*) does not require storing all *k*-mers, the total length of all its edge-labels is up to *k* times smaller than the total length of all edge-labels in the uncompressed de Bruijn graph *UDB*(*Genome*,*k*). The traditional assembly approach constructs *UDB*(*Reads,k*) first and transforms it into *DB*(*Reads,k*). Since this approach is impractical for large genomes and large *k*-mer sizes, jumboDBG directly constructs *DB*(*Reads,k*) without constructing *UDB*(*Reads,k*).

Even though *DB*(*Reads,k*) is more compact than *UDB*(*Reads,k*), its direct construction also requires prohibitively large time/memory. jumboDBG thus assembles reads into disjointigs, sequences that are spelled by arbitrary walks through the (unknown!) graph *DB*(*Reads,k*). Even in the case of error-free reads, a disjointig may represent a misassembled concatenate of segments from various regions of the genome rather than its contiguous substring (Kolmogorov et al., 2019). Although switching from reads to misassembled disjointigs might appear reckless, it is an important step since a carefully chosen disjointig-set *Disjointigs* has a much smaller total disjointig-length than the total read-length while resulting in the same compressed de Bruijn graph *DB*(*Disjointigs,k*) as *DB*(*Reads,k*).

Even though constructing *DB*(*Disjointigs,k*) is an easier task than constructing *DB*(*Reads,k*), it still faces the time/memory bottleneck. jumboDBG addresses it by using the Bloom filter, a compact data structure for storing sets that may report false positives but never false negatives. It stores all (*k*+1)-mers from disjointigs in a Bloom filter formed by multiple independent *hash functions*, each mapping a (*k*+1)-mer into a bit array. The Bloom filter reports a *true positive* for all (*k*+1)-mers occurring in disjointigs but may also report a *false positive* for some (*k*+1)-mers that do not occur in disjointigs (with a small controlled probability). However, the Bloom filter never “forgets” any inserted (*k*+1)-mer and thus never reports a *false negative*.

### Outline of the LJA pipeline

Below we outline all eleven steps of the LJA pipeline. Figure 1 illustrates the work of the jumboDBG module (steps 1-7 of the LJA pipeline). Figure 2 illustrates the entire LJA pipeline. We illustrate the LJA pipeline using the *T2T* dataset of HiFi reads that was semi-manually assembled by the T2T consortium into a sequence *T2TGenome* by integrating information generated by multiple sequencing technologies (CHM13 reference genome version 1.1). All datasets analyzed in this paper are described in Supplementary Note 2.

**Figure 1.**
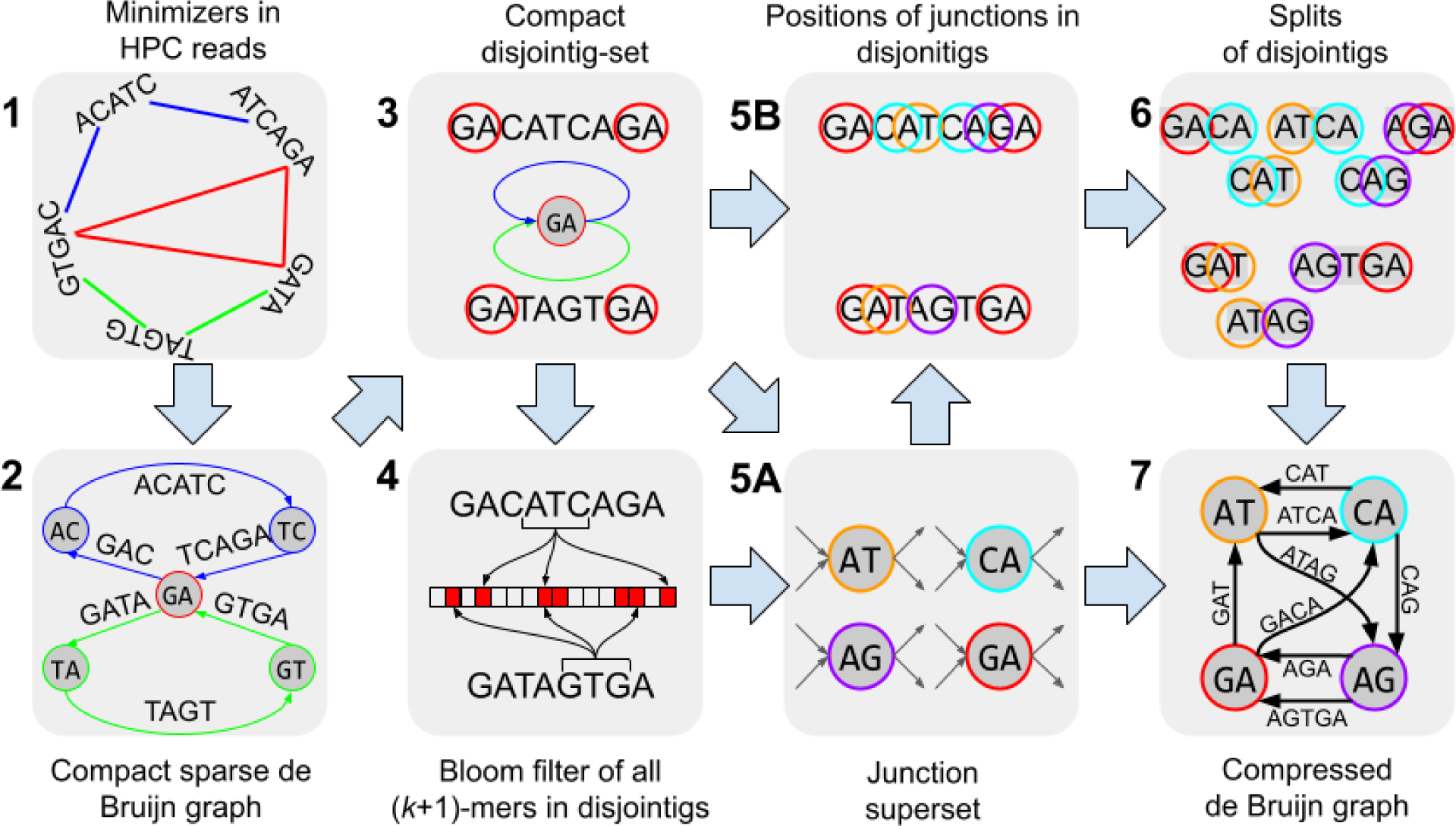
JumboDBG pipeline. (1) Generating the anchor-set *Anchors=*{GA,AC,TC,TA,GT} by finding all minimizers in *Read*s. For simplicity, the figure does not reflect that jumboDBG classifies all *k*-prefixes and *k*-suffixes of reads as minimizers. (2) Constructing a compact sparse de Bruijn graph *SDB*(*Reads,Anchors*). (3) Constructing a compact disjointig-set *Disjointigs* as edge-labels in *SDB*(*Reads,Anchors*). (4) Generating the Bloom filter for all (*k+*1)-mers in disjointigs. Each arrow directed from a (*k+*1)-mer to the Bloom filter illustrates its evaluation by one of the hash functions. (5) Using the Bloom filter to construct the junction-superset *Junctions^+^* and (5B) find positions of *k*-mers from *Junctions^+^* in disjointigs (5A). (6) Breaking disjointigs into splits. (7) Constructing the compressed de Bruijn graph *DB*(*Disjointigs,k*)=*DB*(*Reads,k*).

**Figure 2.**
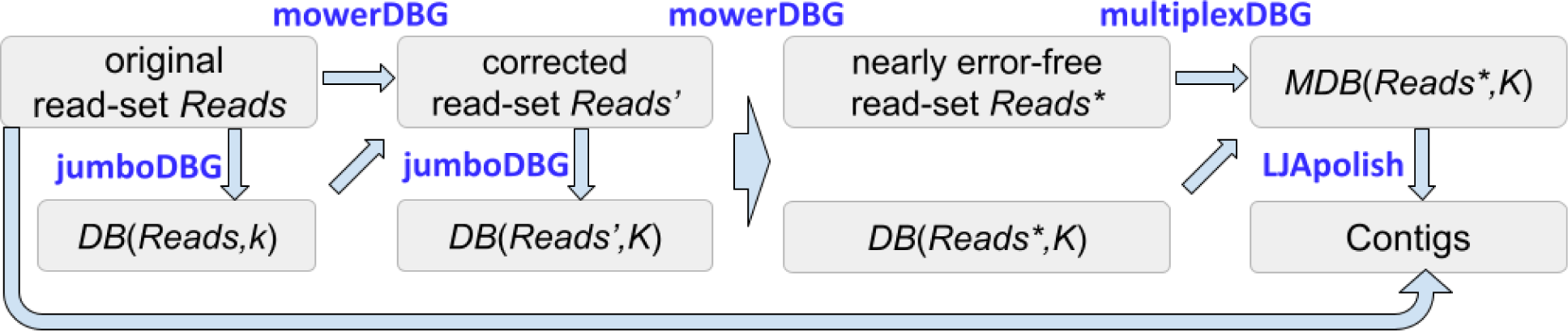
LJA pipeline. jumboDBG first constructs the de Bruijn graph *DB*(*Reads,k*) with a small *k*-mer size. mowerDBG uses this graph to correct errors in reads, resulting in an error-corrected read-set *Reads*’. Afterward, jumboDBG constructs the de Bruijn graph *DB*(*Reads’,K*) on the error-corrected read-set with a large *K-*mer size. mowerDBG uses this graph to correct even more errors in reads, resulting in an error-corrected read-set *Reads*\*. Since error correction in mowerDBG simultaneously modifies the graph *DB*(*Reads’,K*) into the graph *DB*(*Reads*,K*), there is no need to launch jumboDBG again for constructing *DB*(*Reads*,K*). multiplexDBG complements the error-corrected read-set *Reads\** by virtual reads and transforms *DB*(*Reads*,K*) into the multiplex de Bruijn graph *MDB*(*Reads*,K*). LJApolish uses the set of original reads *Reads* to expand HPC contigs formed by non-branching paths in this graph.

### Step 0: Transforming all reads into homopolymer-collapsed (HPC) reads

Since errors in the length of *homopolymer runs* represent the dominant source of errors in HiFi reads (Wenger et al., 2019), LJA collapses each homopolymer run X…X in each read into a single nucleotide X, resulting in a *homopolymer-collapsed* (*HPC*) read. The entire LJA pipeline works with HPC reads, except for the last LJApolish module that expands each collapsed nucleotide X in the resulting HPC assembly into a run X…X of the correct length. On average, uncollapsed reads in the T2T dataset have ≅2000 errors per a megabase of the total read-length. Transforming them into HPC reads reduces the error rate to ≅620 errors per a megabase and makes 38% of all HPC reads error-free when compared to the HPC reference genome.

### Step 1: Generating the anchor-set *Anchors* by finding all minimizers in the HPC read-set *Read*s

Given a hash function defined on *k*-mers and an integer *width*, a *minimizer* of a word of length *width* is defined as a *k*-mer with a minimal hash in this word. A minimizer-set of a string is defined as the set of all minimizers over all its substrings of length *w* (Roberts et al., 2004). We modify the original concept of a minimizer of a linear string by adding its *k*-prefix (prefix of length *k*) and *k*-suffix (suffix of length *k*) to the set of its minimizers. A sensible choice of the parameter *width* (and a hash function) ensures that each read is densely covered by minimizers and that overlapping reads share minimizers, facilitating the assembly. jumboDBG generates the set of all minimizers in reads that we refer to as the *anchor-set Anchors*.

### Step 2: Constructing a compact sparse de Bruijn graph *SDB*(*Reads,Anchors*)

Since the direct construction of *DB*(*Reads,k*) faces the time/memory bottleneck, jumboDBG first assembles reads into disjointigs. Although the Flye assembler (Kolmogorov et al., 2019) constructs disjointings by searching for overlapping reads, it is unclear how to extend this construction to highly-repetitive regions (e.g., centromeres) that Flye does not adequately reconstruct. Instead, jumboDBG constructs a sparse de Bruijn graph and generates disjointigs in this graph that are also disjointigs in the much larger (but unknown) de Bruijn graph.

Given a set of *k*-mers *Anchors* from a string-set *Reads,* we consider each pair *a* and *a*’ of consecutive anchors in each read and generate a substring of the read (called a *split*) that starts at the first nucleotide of *a* and ends at the last nucleotide of *a’*. The resulting set of splits is denoted *Splits*(*Reads,Anchors*). The sparse de Bruijn graph *SDB*(*Reads,Anchors*) is defined as a graph with the vertex-set *Anchors* and the edge-set *Splits*(*Genome, Anchors*). Each string in *Splits*(*Genome, Anchors)* represents the label of an edge connecting its *k*-prefix and *k*-suffix in *SDB*(*Reads,Anchors*).

A sparse de Bruijn graph *SDB*(*Reads,Anchors*) is *compact* if all its vertices (anchors) represent junctions in the graph *DB*(*Reads,k*). jumboDBG transforms the initially constructed graph *SDB*(*Reads,Anchors*) into a compact sparse de Bruijn graph *SDB*(*Reads,Anchors\**).

### Step 3: Constructing a compact disjointig-set *Disjointigs*

A disjointig-set is *complete* if its disjointigs contains all (*k*+1)-mers from *Reads.* A disjointig in the sparse de Bruijn graph *SDB*(*Reads,Anchors*) is *compact* if its *k*-suffix and *k*-prefix are both junctions in the de Bruijn graph *DB*(*Reads,k*). A complete disjointig-set is *compact* if each disjointig in this set is compact. Since *DB*(*Reads,k*)*=DB*(*Disjointigs,k*) for a complete disjointig-set, the problem of constructing the compressed de Bruijn graph from reads is reduced to constructing the compressed de Bruijn graph of a complete disjointig-set. However, not every complete disjointig-set enables efficient construction of this graph. Below we show that a compact disjointig-set enables efficient construction of *DB*(*Disjointigs,k*). jumboDBG constructs a compact disjointig-set as the set of edge-labels in the compact sparse de Bruijn graph *SDB*(*Reads,Anchors\**).

### Step 4: Generating the Bloom filter of all (*k+*1)-mers in disjointigs

Even though we have now reduced constructing the de Bruijn graph *DB*(*Reads,k*) to constructing *DB*(*Disjointigs,k*), even this simpler problem faces the time/memory bottleneck. To address it, jumboDBG constructs the Bloom filter for storing all (*k*+1)-mers in disjointigs and uses the rolling hash to query them in O(1) instead of O(*k*) time.

### Step 5: Using the Bloom filter to construct the junction-superset *Junctions*

The Bloom filter enables rapid construction of a small junction-superset *Junctions^+^* even though the de Bruijn graph of disjointigs has not been constructed yet!

### Step 6. Using the set *Junctions^+^* to break disjointigs into splits

jumboDBG also uses the Bloom filter to rapidly identify the positions of all *k*-mers from *Junctions^+^* in disjointigs and to break disjointigs into splits afterward.

### Step 7. Using splits to construct *DB*(*Disjointigs,k*)=*DB*(*Reads,k*)

An *edge-subpartition* of an edge (*v*,*w*), in a graph “substitutes” it with two edges by “adding” a vertex *u* in the “middle” of this edge. A *subpartition* of a graph is defined as a result of a series of edge-subpartitions. As described in Methods, the string-set *Splits*(*Disjointigs,Junctions^+^*) represents edge-labels of a subpartition of the graph *DB*(*Disjointigs,k*). jumboDBG compresses all 1-in-1-out vertices in this graph to generate *DB(Disjointigs,k*)=*DB*(*Reads,k*).

### Step 8. Correcting errors in reads with mowerDBG and constructing the graph *DB(Reads*,K*) on the error-corrected read-set *Reads\** using a larger *K*-mer size

Supplementary Note 3 illustrates that jumboDBG generates highly contiguous assemblies of a *k*-complete human read-set sampled from *T2TGenome* for large values of *k*, e.g., *k*=5001. However, constructing accurate and contiguous assemblies for real reads is challenging because the de Bruijn graph *DB*(*T2T,k*) of the T2T read-set is much more complex than the de Bruijn graph *DB*(*T2TGenome,k*) even when the error-rate in reads is small. mowerDBG uses the de Bruijn graph *DB*(*Reads,k*) to correct errors in the read-set *Reads.* LJA performs two rounds of error correction and launches mowerDBG twice, with a small *k*-mer size in the first round, resulting in a error-corrected read-set *Reads’*, and a large *K*-mer size in the second round, resulting in a nearly error-free read-set *Reads\** (default values *k*=501 and *K*=5001).

jumboDBG constructs the graph *DB*(*T2T,k*) with 33,230,906 edges in only 2.7 hours using 54 Gb memory footprint. However, over 99% of edges in this graph are triggered by errors in reads: if reads in the T2T dataset were error-free, jumboDBG would construct the de Bruijn graph of the error-free read-set *T2TErrorFree* with only 214,517 edges in 0.6 hours using 33 Gb memory. mowerDBG corrects most errors in reads resulting in a much smaller de Bruijn graph *DB*(*T2T’,k*) with 297,176 edges on the error-corrected read-set *T2T’*. jumboDBG further constructs the compressed de Bruijn *DB*(*T2T’,K*) using a larger *K*-mer size with 79,908 edges. Afterward, mowerDBG performs the second round of error-correction in this graph, resulting in a nearly error-free read-set *Reads\** and a graph *DB(T2T*,K*) with only 6516 edges that approximates the graph *DB(T2TErrorFree,K*) with 4956 edges. We note that since the read-set *T2TErrorFree* was constructed based on the reference *T2TGenome*, that excluded the heterozygous regions present in the CHM13 cell line (Bzikadze and Pevzner, 2020, Nurk et al., 2021), *DB(T2T*,K*) may have some heterozygous edges missing in *DB(T2TErrorFree,K*).

### Step 9: Transforming the de Bruijn graph *DB*(*Reads*,K*) into the multiplex de Bruijn graph

The choice of the *k*-mer size greatly affects the complexity of the de Bruijn graph. There is no perfect choice since gradually increasing *k* leads to a less tangled but more fragmented de Bruijn graph. This trade-off affects the contiguity of assembly, particularly in the case when the *k*-mers coverage by reads is non-uniform, let alone when the read-set misses some genomic *k*-mers. Ideally, we would like to vary the *k*-mer size, reducing it in low-coverage regions (to avoid fragmentation) and increasing it in high-coverage regions (to improve repeat resolution). The *iterative de Bruijn graph* approach (Peng et al., 2010, Bankevich et al., 2012, Peng et al., 2012) is a step toward addressing this goal by incorporating information about the de Bruijn graphs for a range of *k*-mer sizes *k_1_* < *k_2_* < … < *k_t_* into the de Bruijn graph. However, this approach still constructs a graph with a fixed *k_t_*-mer size.

multiplexDBG transforms the compressed de Bruijn graph *DB*(*Reads*,K*) into the multiplex de Bruijn graph *MDB*(*Reads*,K*) with vertices labeled by strings of length varying from *K* to *K^+^*, where *K^+^* is larger than *K* (default value *K*^+^=40,001). It transforms *DB*(*T2T*,*5001) with 6,516 edges into *MDB*(*T2T*,*5001) with only 1432 edges and generates HPC contigs. Note that labels of some vertices of this graph are longer than all reads in the T2T dataset since multiplexDBG adds *virtual reads* to the read-set (Supplementary Note 4).

### Step 10: Expanding HPC contigs

LJApolish expands HPC contigs that represent edge-labels in the multiplex de Bruijn graph and results in an accurate final assembly with only ≅15 single-base errors per million nucleotides.

### Evaluating quality of complete genome assemblies

We used the standard benchmarking metrics (Mikheenko et al., 2018) as well as the additional completeness metric aimed at high-quality HiFi assemblies. We denote the length of a string *s* as |*s*| and the total length of all strings in a string-set *S* as |*S*|. Given a string *s* in a string-set *S*, we define *L^+^*(*s*) as the total length of all strings in *S* with length at least |*s*|. The standard N50 metric for a string-set *S* is defined as the length of the longest string in *S* with *L^+^*(*s*) ≤ 0.5*|*S*|. See Mikheenko et al., 2018 on applying this and similar metrics (e.g., NG50, NGA50, NGA75, or LGA95) to a set of contigs.

We classify a contig as *correctly-assembled* if it has no misassemblies. *Completeness* of a chromosome assembly is defined as the length of the longest correctly-assembled contig from this chromosome divided by the chromosome length (in percentages). A chromosome is *N%*-*assembled* if its completeness is at least *N*%.

As the authors of hifiasm noted (Cheng et al., 2021), the leading assembly evaluation tool QUAST-LG (Mikheenko et al., 2018) has limitations with respect to analyzing misassemblies in highly-repetitive regions. To analyze these limitations, we used QUAST-LG to compare *T2TGenome* with itself. Even though any such comparison should return no errors, QUAST-LG reported 96 global and 222 local misassemblies. The QUAST-LG authors are familiar with this effect, often caused by difficulties in aligning sequences sampled from highly-repetitive regions with long homopolymer runs. Indeed, QUAST-LG reported no global and local misassemblies in comparison of the homopolymer-collapsed *T2TGenome* with itself, suggesting that collapsing homopolymers results in more accurate alignments and thus enables a more accurate analysis of misassemblies. Since QUAST-LG authors recommend using HPC contigs for identifying misassemblies (Dr. Mikheenko, personal communication). we provide information about misassemblies by comparing HPC contigs against HPC references and report the NGA50 and other metrics based on contigs broken at positions defined by misassemblies in HPC contigs.

### Benchmarking LJA

In addition to benchmarking LJA, hifiasm (Cheng et al., 2021) and HiCanu (Nurk et al., 2020) on the T2T dataset, we analyzed their assemblies of human, mouse, maize, and fruit fly HiFi read-sets referred to as HG002, MOUSE, MAIZE, and FLY, respectively. We first benchmark assemblies of the T2T read-set dataset against *T2TGenome* — the only complete and carefully validated large genome reference available today (Mc Cartney et al., 2021).

Table 1 presents benchmarking results for the T2T dataset. LJA produced the most contiguous assembly (NG50=97 Mb) with six 100%-assembled and three additional 95%-assembled chromosomes (compared to NG50=90 Mb and one 95%-assembled chromosome for hifiasm).

**Table 1.**
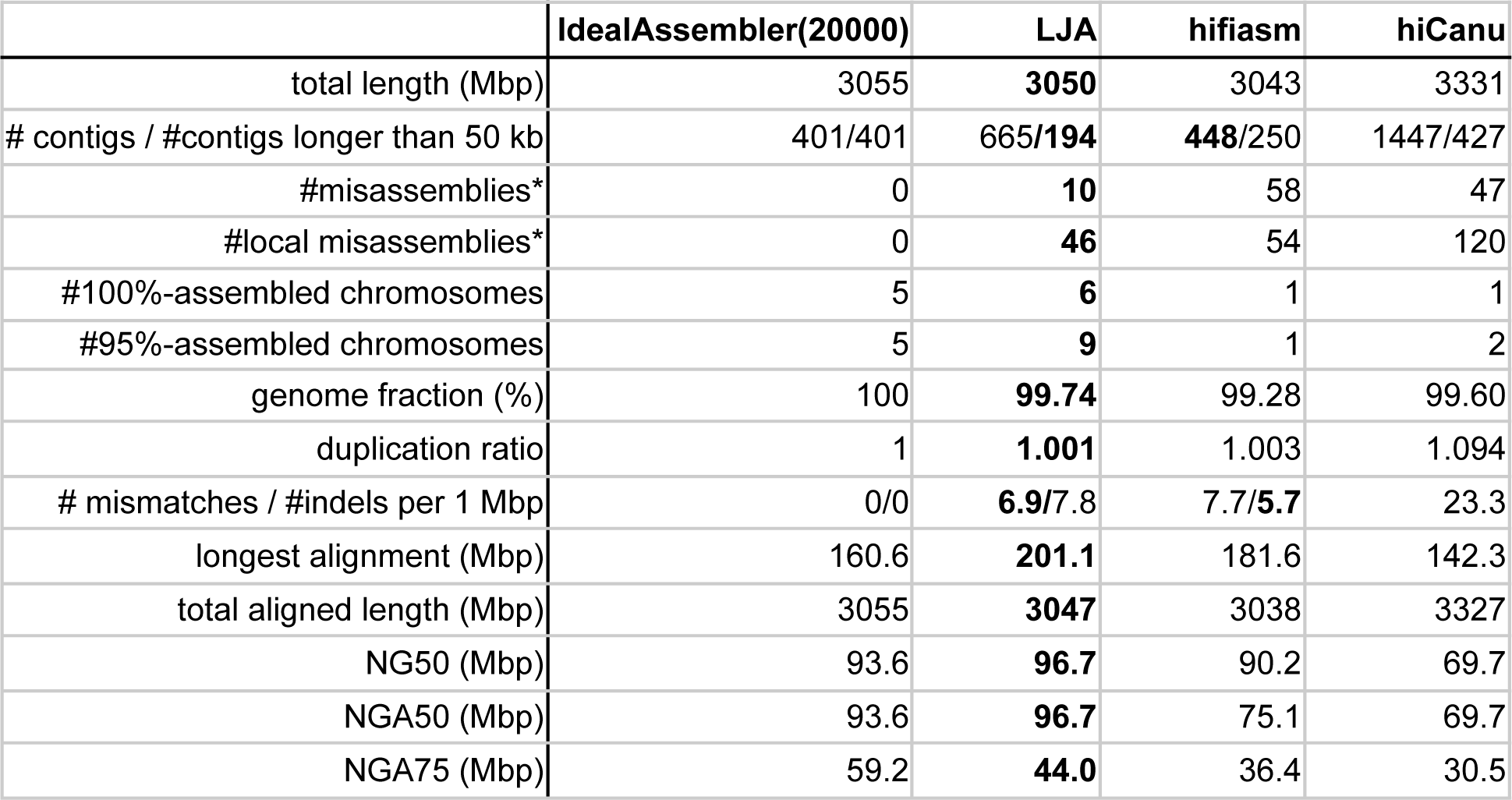
Benchmarking LJA, hifiasm, and HiCanu on the T2T dataset. The assemblies were generated with hifiasm version 0.15.5-r352 and HiCanu version v2.3 and benchmarked using QUAST-LG v5.0.2 with *T2TGenome* as the reference. Misassemblies were identified by a separate run of QUAST-LG with HPC contigs against the HPC reference. LJA 100%-assembled chromosomes 3, 5, 7, 10, 12, and 20, HiCanu 100%-assembled chromosomes 20, and hifiasm 100%-assembled chromosome 5. Since all assemblies may have small variations at the chromosome ends, 99.9%-assembled chromosomes are counted as 100%-assembled chromosomes. “Ideal” refers to the theoretically optimal assembly of a *k*-complete read-set obtained by generating contigs as edge-labels of the graph DB(*T2TGenome*,20000). Note that, even though very few HPC reads in the T2T dataset are longer than 20000 bp, the LJA assembly of this read-set may improve on the ideal assembly DB(*T2TGenome*,20000) because it utilizes information about coverage for loop resolution (See Supplementary Note 5). The reference length is 3,054,832,041. The number of LJA contigs (665) is smaller than the number of edges in the constructed multiplex de Bruijn graph (1432) because removing overlaps between contigs in the final LJA output results in many short contigs (LJA removes contigs that become shorter than 5 kb).

We classify a contig with length larger or equal than NG50 (NG75) as *NG50*-*long (NG75*-*long)* contig. It turned out that 1 out of all 12 NG50-long contigs assembled by LJA and 4 out of all 12 NG50-long contigs assembled by hifiasm are misassembled, resulting in a reduced NGA50=81 Mb for hifiasm. LJA misassembled only 1 out of its 24 NG75-long contigs while hifiasm misassembled 9 out of its 23 NG75-long contigs. Figure 3 demonstrates that LJA significantly improves on both hifiasm and HiCanu with respect to chromosome-by-chromosome and centromere-by-centromere assemblies.

**Figure 3.**
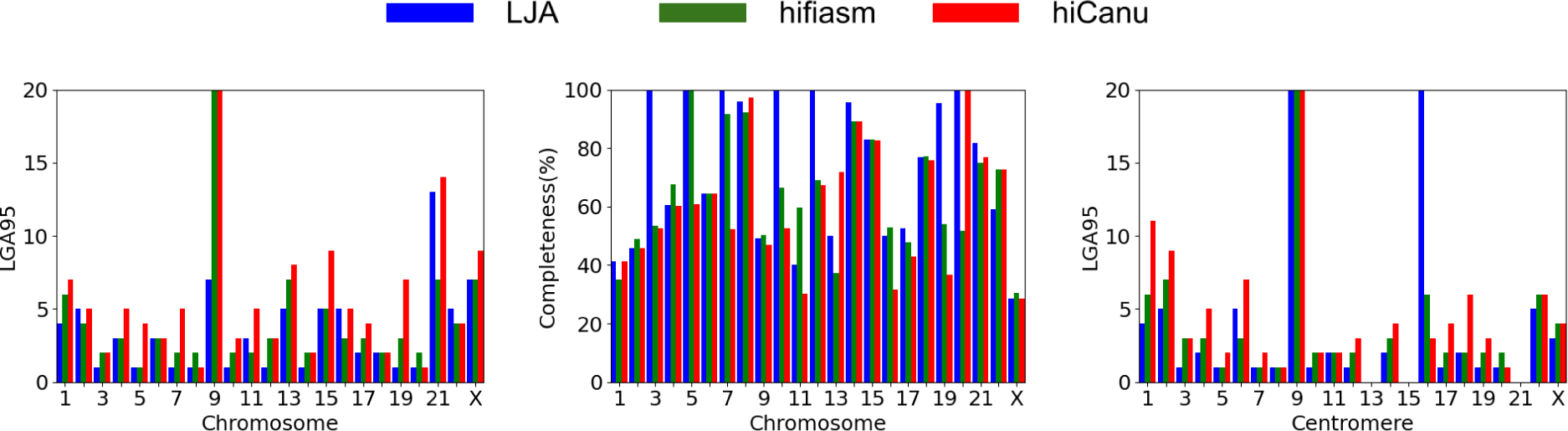
Chromosome-by chromosome LGA95 (left) and completeness (center) metrics as well as centromere-by-centromere LGA95 metric (right) for LJA (blue), hifiasm (green), and HiCanu (red) assemblies of the T2T read-set. LGA95 for each chromosome (centromere) is defined as the number of contigs (fragments of contigs broken at misassembly breakpoints) needed to cover 95% of its length. LJA 100%-assembled six chromosomes and resulted in a better assembly (e.g., assembly with lower LGA95) than hifiasm and HiCanu for most chromosomes. Although hifiasm resulted in lower LGA95 than LJA on chromosomes 2 (4 vs 5), 11 (2 vs 3), 16 (3 vs 5), 21 (7 vs 13), and 22 (4 vs 5), it made 10 misassemblies in these five chromosomes compared to only 1 misassembly made by LJA. For example, for chromosome 21 with the biggest gap in the LGA95 values between hifiasm and LJA (7 vs 13), LJA resulted in a higher completeness (82%) than hifiasm (75%). As another example, LJA resulted in a much larger LGA95=22 on centromere 16 than hifiasm (6) and HiCanu (3) but made no errors (both hifiasm and HiCanu had 3 misassemblies). LGA95 values for centromeres 13, 15 and 21 (for all assemblers) are undefined because QUAST-LG reports multiple gaps in these centromeres. Long and highly repetitive rDDNA arrays are the only regions in the human genome that were not assembled by the T2T consortium. These regions are represented as simulated *rDDNA models* rather than correct rDDNA sequences in *T2TGenome*. Since centromeres 13, 15 and 21 include rDDNA models, QUAST-LG reports multiple gaps in their coverage by contigs that sum up to more than 5% of the lengths of these centromeres.

LJA significantly reduced the number of misassemblies (10) as compared to hifiasm (58) and HiCanu (47). Analysis of misassemblies reported by QUAST-LG in 30 longest contigs (for each assembly tool) using the Icarus visualization tool (Mikheenko et al., 2016) confirmed that they all represent structural errors (a large insertion, deletion or relocation) rather than alignment artifacts. While LJA produced more contigs than hifiasm (665 vs 448), this increased number does not indicate an inferior assembly but rather reflects the fact that LJA assembled an extra 13 Mb of the genome as compared to hifiasm (99.7% vs. 99.3% genome fraction). In fact, hifiasm produced more contigs longer than 50 Kb than LJA (250 vs 194).

LJA and hifiasm made an order of magnitude fewer single-base errors (mismatches and indels) than HiCanu. Analysis of these errors should take into account that an assembler covering a larger fraction of a genome may have additional errors in highly-repetitive regions that are not covered by other assemblers. Although LJA made slightly more errors than hifiasm (34293 vs. 33732), it made a smaller number of errors in regions assembled by both LJA and hifiasm (2330 LJA errors were made in highly-repetitive regions that were not assembled by hifiasm).

Table 1 also benchmarks the “ideal” assembler on a *k*-complete read-set that outputs contigs as the edge-labels of the graph DB(*T2TGenome*,20000). It turned out that LJA assembly is similar in quality to the theoretically optimal “ideal” assembly of error-free reads of length 20000.

### Assembling a diploid human genome

Both short-read and long-read assemblers often collapse two heterozygous alleles (typically represented as bulges in the de Bruijn graph) into a single copy to increase the contiguity of the *consensus assembly*, a mosaic of segments from maternal and paternal chromosomes. Diploid assemblers try to prevent such collapsing and include two steps: (i) constructing a *phased assembly graph* that accurately represents the heterozygous alleles, and (ii) using the entire read-lengths and complementary technologies (ultralong reads, Hi-C reads, etc.) to increase the contiguity of paternal/maternal contigs in the phased assembly graph and generate a *haplotype-resolved assembly* (Cheng et al., 2021, Garg et al., 2021). Constructing an accurate and contiguous phased assembly graph is a critical step in both consensus and diploid assembly and the focus of the current efforts of the T2T Project as it scales up from a single haploid to multiple diploid genomes. Indeed, a fragmented phased graph makes it difficult to generate both consensus and haplotype-resolved assemblies, not to mention that each error in the graph likely triggers downstream errors in these assemblies.

LJA does not collapse heterozygous alleles and thus automatically generates a phased de Bruijn graph of a diploid genome. Although generating the consensus and haplotype-resolved assemblies are outside the scope of this paper, the phased LJA graph represents an excellent starting point for generating these assemblies.

We analyzed the phased LJA and hifiasm assemblies using the HG002 read-set from a diploid human genome. LJA generated an assembly with N50=383 kb and total length 5.8 Gb, while hifiasm generated an assembly with N50=310 kb and total length 6.7 Gb (before heterozygosity collapsing) that is ≅2.2 times larger than the human genome length. While LJA generated a more contiguous phased assembly than hifiasm, it is unclear how to evaluate the accuracy of these assemblies in the absence of a validated complete reference genome for each haplome.

Although the LJA graph of the diploid read-set is much larger than the graph of the haploid T2T read-set (42298 vs 1440 edges), it has a rather simple “bulged” structure, making it well-suited for the follow-up consensus and haplotype-resolution steps. We define the *bulge index* of a graph as the fraction of edges that form bulges in this graph (an edge forms a bulge if it has a parallel edge). LJA constructs the multiplex de Bruijn graph with the bulge index 85% for the HG002 read-set, illustrating that most of the graph is formed by bulges. Each bulge represents differing segments of paternal and maternal alleles with average length ≅150 kb alternating with much shorter unique regions (identical alleles with average length ≅30 kb).

### Assembling mouse, maize, and fruit fly genomes

We benchmarked LJA, hifiasm, and HiCanu on the MOUSE, MAIZE, and FLY read-sets from inbred species. Although the reference genomes for these datasets are available, it is unclear how to compare them with HiFi assemblies since the quality of these assemblies may exceed the quality of the references. For example, while Table 2 illustrates that LJA and hifiasm improved on HiCanu with respect to NG50 metric, this metric does not account for errors in assemblies and errors in references. E.g., hifiasm assembled the MAIZE dataset with the highest NG50 (35 Mb vs 26 Mb for LJA) but made more misassemblies (6081 vs 5594 for LJA), resulting in a rather low NGA50 (1.4 Mb for both hifiasm and LJA). The extremely large number of misassemblies for the MAIZE dataset suggests that many of them may be triggered by errors in the reference genome or differences between maternal and paternal alleles (even after inbreeding).

**Table 2.** Benchmarking LJA, hifiasm, and HiCanu on MOUSE, MAIZE, and FLY read-sets. Mouse, maize, and fruit fly samples represent the C57BL/6J strain of *Mus musculus* (Hon et al., 2020), the B73 strain of *Zea mays* (Hon et al., 2020), and *Drosophila ananassae* (Tvedte et al., 2021), respectively. We used estimates of the lengths of the mouse, maize and fly genomes (2.7 Gb, 2.3 Gb, and 0.22 Gb, respectively) for NG50 calculation

| assembler | MOUSE |  |  | MAIZE |  |  | FLY |  |  |
| --- | --- | --- | --- | --- | --- | --- | --- | --- | --- |
|  | #contigs | total length (Gb) | NG50 (Mb) | #contigs | total length (Gb) | NG50 (Mb) | #contigs | total length (Gb) | NG50 (Mb) |
| LJA | 1282 | 2.71 | 24 | 1310 | 2.15 | 26 | 313 | 0.22 | 9.5 |
| hifiasm | 658 | 2.61 | 21 | 1136 | 2.18 | 35 | 933 | 0.24 | 10.0 |
| HiCanu | 3334 | 2.67 | 15 | 1992 | 2.17 | 23 | 6439 | 0.32 | 9.2 |

The fruit fly *D. ananassae* genome, first assembled in 2007 and recently reassembled using five different technologies, represents one of the most contiguous and accurate eukaryotic assemblies available today (Miller et al., 2018, Tverdte et al, 2021). However, QUAST-LG reported two orders of magnitude more LJA misassemblies (625) for the comparatively small fly genome than for the entire *T2TGenome* even after homopolymer collapsing (964 misassemblies for hifiasm and 1082 misassemblies for HiCanu). Although the LJA assembly is much closer to the reference than the hifiasm and HiCanu assemblies, the unusually high number of misassemblies suggests that many of them may be triggered by errors in the reference genome or differences between maternal and paternal alleles. Another indication of potential single-base errors in the reference is the greatly increased rate of single-nucleotide errors as compared to the T2T dataset (527, 599, and 926 errors per megabase for LJA, hifiasm, and HiCanu assemblies, respectively). Also, while the *D. ananassae* genome is sequenced from an inbred organism, the coverage of *k*-mers in reads reveals significant heterozygosity (Supplementary Note 5) that makes the assembly quality evaluation challenging.

Our analysis suggests that, despite multiple previous attempts to assemble the *D. ananassae* reference genome, it likely has many misassemblies. It also illustrates that existing HiFi assemblers generate rather different results (even in a relatively simple case of inbred genomes), implying that at least some of them make many assembly errors. In the absence of an automated assembly validation pipeline (Mc Cartney et al., 2021), It remains unclear how to detect these errors (even with using complementary technologies) and generate error-free references in the era of complete genome sequencing.

### Benchmarking jumboDB

Supplementary Note 3 provides information about benchmarking individual modules of the LJA pipeline. Below we focus on benchmark the jumboDBG module that defines the memory footprint and accounts for roughly one third of the running time of the entire LJA pipeline

Comparison of two subfigures in Figure 4 illustrates that the running time and memory footprint of jumboDBG are largely defined by the size of the constructed de Bruijn graph rather than the *k*-mer size. When the *k*-mer size increases from 501 to 5001, the running time of jumboDBG on the T2T read-set increases from 2.7 hours to 3.9 hours (memory increases from 54 Gb to 93 Gb), while the running time (≅0.6 hours) and memory (≅35 Gb) on the T2TErrorFree read-set remains roughly the same. We did not benchmark jumboDBG against other tools since there are no tools for constructing large compressed de Bruijn graphs yet: the sparse de Bruijn graph constructed by MBG (Rautiainen and Marschall, 2021) and repeat graph constructed by Flye (Kolmogorov et al., 2019) represent approximations of the compressed de Bruijn graph.

**Figure 4.**
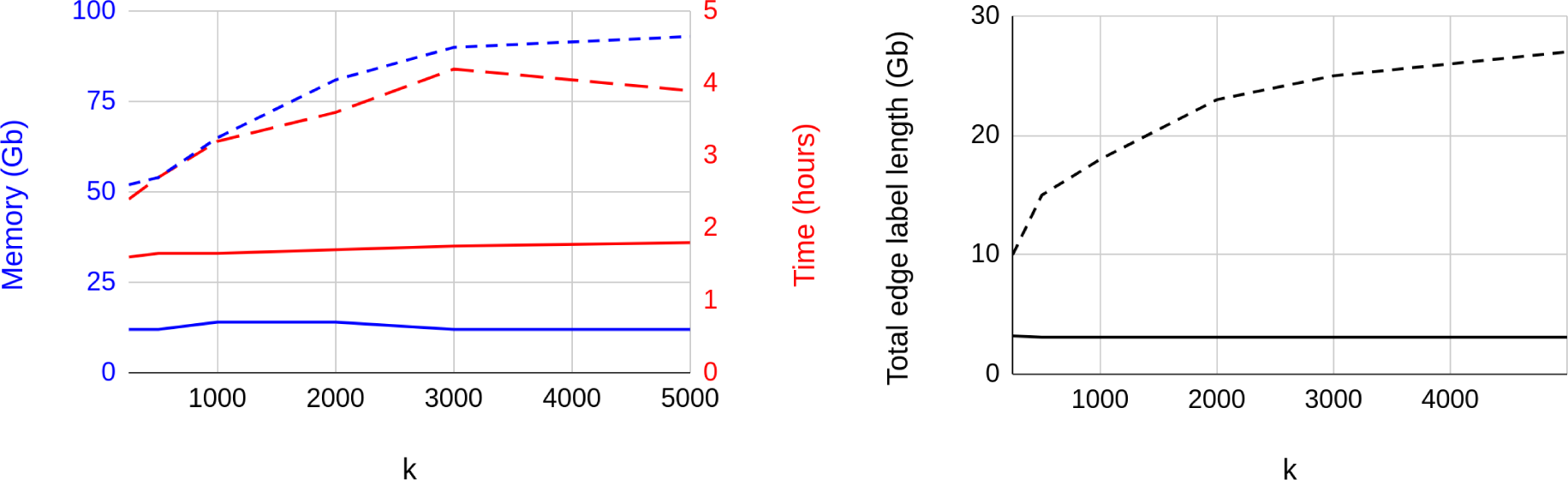
The running time and memory footprint of jumboDB. (Left) The running time (red) and memory footprint (blue) of jumboDBG on the T2T read-set (dashed lines) and the T2TErrorFree read-set (solid lines) for *k-*mer sizes ranging from 251 to 5001. (Right) The total length of all edges in the de Bruijn graphs *DB*(*T2T*,*k*).

## Methods

The Methods section is organized as follows. Before constructing the compressed de Bruijn graph *DB*(*Reads,k*), we describe a simpler yet still open problem of constructing the compressed de Bruijn graph *DB*(*Genome*,*k*) of a large *circular* string *Genome* for a large *k*-mer size. This problem (Minkin et al., 2017) serves as a stepping stone for constructing a much larger graph *DB*(*Reads,k*). After describing the algorithm for constructing the compressed de Bruijn graph of a circular genome, we describe complications that arise in the case of constructing the compressed de Bruijn graph from a genome with linear chromosomes. Afterwards, we describe how to modify the algorithm for constructing *DB*(*Genome,k*) into the algorithm for constructing *DB*(*Reads,k*). Since this transformation faces the time/memory bottleneck, we describe how jumboDBG first assembles reads into disjointigs. Afterwards, we describe steps 8 (error-correcting reads), 9 (constructing multiplex de Bruijn graph), and 10 (expanding HPC assembly) of the LJA pipeline. Supplementary Note 6 describes LJA parameters.

Since the algorithm for constructing the de Bruijn graph of a circular genome does not require construction of disjointigs (step 3), we number its steps 4-7 as 4G-7G in a way that is consistent with the previously described steps of jumboDB:

**Step 4G:** Generating the Bloom filter for all (*k+*1)-mers in *Genome*.

**Step 5G:** Using the Bloom filter to construct a junction-superset *Junctions^+^* of *Genome*.

**Step 6G:** Using the set *Junctions^+^* to break *Genome* into splits.

**Step 7G:** Using splits to construct *DB(Genome,k*).

### Sparse de Bruijn graphs

Given a set of *k*-mers *Anchors* from a string-set *Genome,* the sparse de Bruijn graph *SDB(Genome,Anchors*) is a graph with the vertex-set *Anchors* and the edge-set *Splits*(*Genome, Anchors*) (each split in *Splits*(*Genome, Anchors)* represents a label of an edge connecting its *k*-prefix with its *k*-suffix). Two vertices in this graph may be connected by multiple edges corresponding to different splits with the same *k*-prefixes and *k*-suffixes (edges labeled by the same sequences are glued together). A straightforward algorithm for constructing *SDB*(*Genome,Anchors*) takes O(|*Genome*|・|*Anchors*|・*k*) time.

A string-set *Genome* corresponds to a path-set that traverses each edge in the de Bruijn *DB*(*Genome,k*) at least once and spells *Genome*. We refer to this path-set as the *genome traversal*. When the anchor-set coincides with the junction-set *Junctions*=*Junctions*(*Genome,k*), each vertex of *DB*(*Genome,k*) is an anchor and each edge corresponds to two consecutive anchors in the genome traversal. Therefore, the sparse de Bruijn graph *SDB*(*Genome,Junctions*) coincides with *DB*(*Genome,k*). Moreover, if *Junctions^+^* is a *superset* of all junctions, that contains all junctions as well as some *false junctions* (i.e., non-branching *k*-mers from *Genome*), *SDB*(*Genome,Junctions^+^*) is a subpartition of *DB*(*Genome,k*).

The key observation is that if the junction-set *Junctions=Junctions*(*Genome,k*) was known, construction of *DB*(*Genome,k*) would be a simple task because it coincides with *SDB*(*Genome,Junctions*). Moreover, even if the junction-set is unknown but a junction-superset *Junctions^+^* (with a small number of false junctions) is known, one can construct *DB*(*Genome,k*) by constructing *SDB*(*Genome,Junctions^+^*) and compressing all its non-branching paths.

### Step 4G: Generating the Bloom filter for all (*k+*1)-mers in *Genome*

In the case of assembling reads, jumboDBG stores all (*k*+1)-mers from disjointigs in a Bloom filter *Bloom*(*Disjointigs,k,BloomNumber,BloomSize*) formed by *BloomNumber* different independent *hash functions*, each mapping a (*k*+1)-mer into a bit array of size *BloomSize* (in the case of a circular genome, it assumes that this genome forms a single disjointig). Storing all (*k*+1)-mers in a Bloom filter allows one to quickly query whether an arbitrary (*k*+1)-mer occurs in dosjointigs and thus speed-up the de Bruijn graph construction (Pell et al., 2012, Chikhi and Rizk, 2013). Given hash functions *h_1_,h_2_,…,h_BloomNumber_* and a *k*-mer *a*, one can quickly check whether all bits *h_1_(a), h_2_(a),…h_BloomNumber_* of the Bloom filter are equal to 1, an indication that the *k*-mer *a* may have been stored in the Bloom filter. Supplementary Note 6 describes how jumboDBG sets the parameters of the Bloom filter.

### Step 5G: Using the Bloom filter to construct the junction-superset *Junctions^+^* of *Genome*

To generate a junction-superset *Junctions^+^* with a small number of false junctions, jumboDBG uses the Bloom filter to compute the upper bound on the indegree and outdegree of each vertex in *UDB(Genome,k*) (a *k*-mer from *Genome*) by checking which of its 4+4=8 forward and backward extensions by a single nucleotide represent (*k*+1)-mers present in the Bloom filter. A *k*-mer is called a *joint* if the upper bounds on either its indegree or outdegree exceed 1. Since each junction is a joint, the set of all joints forms a junction-superset.

A vertex in a graph is *complex* if both its indegree and outdegree exceed 1, and *simple*, otherwise. A junction is a *dead-end* if it has no incoming or no outgoing edges, and a *crossroad,* otherwise. In the case of linear chromosomes, the Bloom filter may overestimate the indegree and/or outdegree of some dead-end junctions, e.g., to misclassify a 0-in-1-out junction as a simple vertex. However, all crossroad junctions will be correctly identified, thus solving the problem of generating a junction-superset (in the case of a circular genome that does not have dead-end junctions), constructing a subpartition of *DB*(*Genome,k*), and further transforming this subpartition into *DB*(*Genome,k*).

### Steps 6G and 7G: Using the set *Junctions^+^* to break *Genome* into splits and using splits to construct *DB*(*Genome*,*k*)

As described above, to construct *SDB*(*Genome,Junctions^+^*), one can generate a Bloom filter for computing a junction-superset *Junctions^+^* and check whether each *k*-mer from *Genome* coincides with a *k*-mer from *Junctions^+^*. Both these tasks require computing hashes of each *k*-mer, a procedure that usually takes O(*k*) time and becomes slow when *k* is large. For example, constructing a hashmap of *Junctions^+^* results in a prohibitively slow algorithm for constructing *SDB*(*Genome,Junctions^+^*) with *O(|Genome|*・*k*) running time.

To compute the hash function in O(1) rather than O(*k*) time, jumboDBG uses a 128-bit polynomial rolling hash (Karp and Rabin, 1987) of *k*-mers from the genome to rapidly check whether two *k*-mers (one from *Genome* and one from *Junctions^+^*) are equal and to reduce the time to construct the compressed de Bruijn graph to *O(|Genome|)*. Similarly, to speed-up the construction of *Junctions^+^*, instead of storing *k*-mers, jumboDBG stores their rolling hashes in the Bloom filter, thus reducing the running time from O(|*Genome*|・*k*) to *O(|Genome|*). Therefore, it constructs the compressed de Bruijn graph of a circular genome in *O(|Genome|*) time that does not depend on the *k*-mer size.

### The challenge of constructing the compressed de Bruijn graph from reads

The described algorithm for constructing the de Bruijn graph of a circular genome can be applied to any string-set *Genome*, resulting in a graph that we refer to as *DB\**(*Genome,k*). However, although *DB\**(*Genome,k*)=*DB*(*Genome,k*) for a genome formed by circular chromosomes, it is not the case for a genome with linear chromosomes (or a genome represented by a *k*-complete error-free read-set).

We say that a string-set *Genome bridges* an edge of the compressed de Bruijn graph *DB*(*Genome,k*) if the label of this edge represents a substring of *Genome*. A genome is called *bridging* (with respect to a given *k*-mer size) if it bridges all edges of *DB*(*Genome,k*), and *non-bridging* otherwise. For example, a *Genome* formed by “chromosomes” ATGC and GCACC is non-bridging since *DB*(*Genome,*2) consists of a single edge with label ATGCACC that does not represent a substring of *Genome*.

Although *DB\**(*Genome,k*)=*DB*(*Genome,k*) in the case of a bridging genome, *DB*(Genome,k*) does not necessarily coincide with *DB*(*Genome, k*) for a non-bridging genome, e.g., a genome with linear chromosomes that does not bridge all edges of *DB*(*Genome,k*). However, after extending the junction-superset by *k*-prefixes and *k*-suffixes of all linear chromosomes, the same algorithm will construct the graph that represents a subpartition of *DB*(*Genome,k*). Although this subpartition can be further transformed into *DB*(*Genome,k*) by compressing all non-branching paths, the resulting algorithm becomes slow when the number of linear chromosomes is large, resulting in a prohibitively large increase in the size of the junction-superset. This increase becomes problematic when one constructs the compressed de Bruijn graph *DB*(*Reads, k*) since each read represents a linear “mini-chromosome”. Even more problematic is the accompanying increase in the number of calls to the hash functions that scales proportionally to the coverage of the genome by reads. An additional difficulty is that, in the absence of the genome, it is unclear how to select the appropriate size of the Bloom filter that keeps the false-positive rate low — selecting it to be proportional to the total read-length (as described in Supplementary Note 6) results in the prohibitively large memory footprint. Even if the genome size is known it is unclear how to select *BloomSize* since the number of different *k*-mers in reads affects the false-positive rate.

jumboDBG addresses these problems by assembling reads into compact disjointigs that form a bridging genome for the graph *DB*(*Reads,k*) and constructing the compressed de Bruijn graph *DB*(*Disjointigs,k*)=*DB*(*Reads,k*) from the resulting disjointig-set *Disjointigs* instead of the read-set *Reads*. Using disjointigs allows us to set the parameter *BloomSize* to be proportional to the total disjointig-length (that is typically an order of magnitude smaller than the total read-length), thus greatly reducing the memory footprint.

### Step 3: Constructing a compact disjointig-set from a read-set

We defined the concepts of complete and compact disjointig-sets in the Results section. Similarly to a disjointig of a read-set, a disjointig of a genome is defined as a string spelled by an arbitrary path in *DB*(*Genome,k*). If a disjointig-set *Disjointigs* is complete then *DB*(*Genome,k*) *= DB*(*Disjointigs,k*). However, the graph *DB\**(*Disjointigs,k*) constructed by jumboDBG may differ from *DB*(*Genome,k*) since it does not include edges of *DB*(*Genome,k*) that are not bridged by *Disjointigs*. However, if a disjointig-set *Disjointigs* is compact, it forms a bridging genome, implying that *DB\**(*Disjointigs,k*) = *DB*(*Genome,k*).

jumboDBG constructs a compact disjointig-set as a set of edge-labels in the compact sparse de Bruijn graph *SDB*(*Reads,Anchors\**). Traditionally the anchor-set *Anchors* for constructing *SDB*(*Reads,Anchors*) is constructed as a set of all minimizers across all reads. However, if the *k*-prefix and/or the *k*-suffix of a read are not anchors, they may be missing in *SDB*(*Reads,Anchors*) since only segments between anchors are added to this graph. As described in Results, jumboDBG modifies the concept of a minimizer of a linear string by adding its *k*-prefix and *k*-suffix to the set of its minimizers and resulting in the set *Anchors=Anchors*(*Reads,width,k*). jumboDBG constructs the sparse de Bruijn graph *SDB*(*Reads,Anchors*), transforms it into a compact sparse de Bruijn graph *SDB*(*Reads,Anchors\**) as described in Supplementary Note 7, and generates a compact disjointig-set as labels of all non-branching paths in this graph.

### Step 8. Correcting errors in reads and constructing the graph *DB(Reads*,K*) on the error-corrected read-set *Reads\** using a larger *K*-mer size

Since an error in a single position of a read triggers an error in each *k*-mer covering this position and since a typical HiFi read has one error per 500 nucleotides on average, the fraction of correct *k*-mers (among all *k*-mers in reads) becomes rather low when *k* exceeds 501, resulting in a complex de Bruijn graph of reads that does not adequately represent the genome. Ideally, we would like to construct the de Bruijn graph using as large *k* as possible, e.g., *k*=15000, slightly below the typical read-length in the T2T dataset. However, this approach results in a highly fragmented de Bruijn graph since reads in this dataset do not span a large fraction of genomic 15000-mers. Although reducing *k* to, lets say, 5000 addresses this complication (nearly all genomic 5000-mers are spanned by reads), most 5000-mers in reads are erroneous, preventing their assembly.

LJA attempts to minimize the effect of (i) errors in reads and (ii) incomplete coverage of genomic *k*-mers by constructing and error-correcting the de Bruijn graphs with a small *k*-mer size and a large *K*-mer size with default values *k*=501 and *K*=5001. As described in Results, this two-round error-correction procedure results in an nearly error-free read-set *Reads*\* (Figure 2).

mowerDBG utilizes the de Bruijn graph of HPC reads for detecting errors in these reads. Since ≅69% of 501-mers in HPC reads are correct, the correct 501-mers have high coverage, in contrast to low-coverage erroneous 501-mers that form *bulges* and *tips* (with the exception of *k*-mers that contain some tandem dinucleotide repeats discussed in Supplementary Note 8). Afterward, mowerDBG uses the *path-rerouting* and *bulge-collapsing* error-correction approaches to simultaneously correct reads and the graph.

Even though the previous paragraph might create an impression that errors in long and accurate reads can be corrected by simply applying error-correcting approaches developed for short reads (e.g. from SPAdes assembler), correction of HiFi reads faces unique challenges that we outline below. First, our goal is to correct reads (rather than to correct the de Bruijn graph as in SPAdes) since we need to rescue as many correct *k*-mers as possible for the second round of error correction with a larger *K*-mer size. Second, we need to perform nearly perfect error correction even in highly-repetitive regions (e.g., centromeres) that short-read assemblers like SPAdes do not even try to assemble. Third, in difference from the short-read assembly, the target *k*-mer size (501) is a small fraction of a typical read-length (15000). As a result, analyzing a bulge in a highly-repetitive region requires not only analysis of the graph structure (like in SPAdes) but also analyzing all read-paths traversing this bulge.

To address these complications, mowerDBG complements path-rerouting and bulge-collapsing by additional steps referred to as correcting dimers and correcting pseudo-correct reads. Supplementary Note 8 describes the mowerDBG algorithm.

### Step 9: Transforming the de Bruijn graph into the multiplex de Bruijn graph

Below we describe a graph transformation algorithm for transforming the graph *DB*(*Reads,k*) into *DB*(*Reads,k^+^*) for *k^+^*>*k* by iteratively increasing the *k*-mer size by 1 at each iteration. Although launching jumboDBG to construct *DB*(*Reads,k*), followed by these transformations, takes more time than simply launching jumboDBG to construct *DB*(*Reads,k^+^*), we use it as a stepping stone toward the multiplex de Bruijn graph construction.

Below we consider graphs, where each edge is labeled by a string, and each vertex *w* is assigned an integer v*ertexSize*(*w*) ≥ *k.* We limit attention to graphs where suffixes of length *vertexSize*(*w*) for all incoming edges into *w* coincide with prefixes of length *vertexSize(w)* for all outgoing edges from *w*. We refer to the string of length *vertexSize*(*w*) that represents these prefixes/suffixes as the label of the vertex *w*. Below we consider graphs with specified edge-labels (vertex-labels can be inferred from these edge-labels) and assume that different vertices have different vertex-labels. We will start by analyzing graphs with the same vertex size for all vertices and will later transition to the multiplex de Bruijn graphs that have vertices of varying sizes.

#### Transition-graph

Let *Transitions* be an arbitrary set of pairs of consecutive edges (*v,w*) and (*w,u*) in an edge-labeled graph *G*. We define the *transition-graph G*(*Transitions*) as follows. Every edge *e* in *G* corresponds to two vertices *e_start_* and *e_end_* in *G*(*Transitions*) that are connected by a blue edge that inherits the label of the edge *e* in *G* (Figure 5.A) We set *vertexSize*(*e_start_*)*=vertexSize*(*e_end_)=k+*1 (vertex-labels are uniquely defined by the (*k*+1)-suffixes/prefixes of the incoming/outgoing edges in each vertex). If an edge *e* in *G* is labeled by a (*k+*1)-mer, the corresponding blue edge in *G*(*Transitions*) is collapsed into a single vertex *e_start_*=*e_end_*. In addition to blue edges, each pair of edges *in=*(*v,w*) and *out*=(*w,u*) in *Transitions* adds a red *transition edge* between *in_end_* and *out_start_* to the transition graph. The label of this red edge is defined as a (*k*+2)-mer *symbol_-(k+1)_*(*in*)*\*label*(*w*)*\*symbol_(k+1)_*(*out*), where *symbol_i_*(*e*) stands for the *i*-th symbol of *label*(*e*) and *symbol_-i_*(*e*) stands for the *i*-th symbol from the end of *label*(*e*).

**Figure 5.**
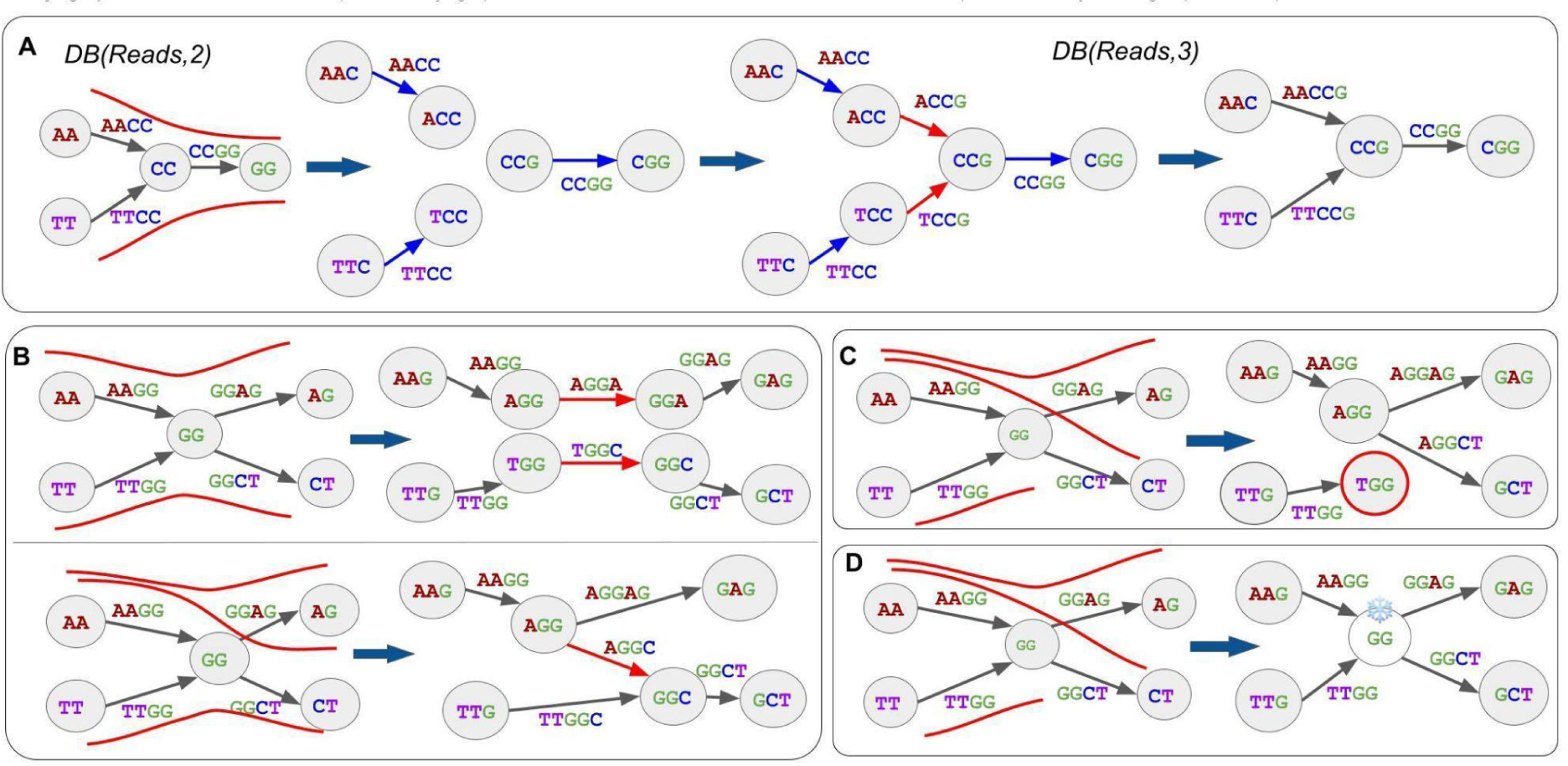
Constructing multiplex de Bruijn graph. **(A) Iterative construction of the compressed de Bruijn graph.** Transformation of *DB*(*Reads*,2) into *DB*(*Reads*,3) for the read-set *Reads=*{**AA`CC`GG**,**TT`CC`GG**}. The first read defines a transition between edges **(AA,CC)** and **(CC,GG)** while the second read defines a transition between edges **(TT,CC)** and **(CC,GG)**. The resulting transition-set defines a transition-graph that coincides with *DB*(*Reads*,3). The read-paths that define the transition-set are depicted as red curves traversing *DB*(*Reads*,2). multiplexDBG “tears apart” edges of the *DB*(*Reads*,2) and increases the *k*-mer length within the vertices of the resulting isolated edges. Afterward, it introduces new red edges that correspond to connections induced by transitions, resulting in a path-graph. Compressing non-branching paths in the path-graph (ignoring the edge colors) results in *DB*(*Reads*,3). Since vertex “**CC”** is simple, graphs *DB*(*Reads*,2) and *DB*(*Reads*,3) have the same topology. The shown transformation merely substitutes the *k*-mer label of this vertex by the (*k*+1)-mer *label*(*w*)*\*symbol_k+1_*(*out*). It preserves the label of the outgoing edge from this vertex and adds a single symbol to the labels of incoming edges into this vertex. **(B) Transforming a complex vertex.** The read-sets *Reads*_1_={**AAGGAG,TTGGCT} (**above) and *Reads*_2_={**AAGGAG, TTGGCT, AAGGCT} (**below) result in the same graphs *DB*(*Reads*_1_,2) and *DB*(*Reads*_2_,2) but different graphs *DB*(*Reads*_1_,3) and *DB*(*Reads*_2_,3). Since vertex **GG** in the graph *DB*(*Reads*_1_,2)=*DB*(*Reads*_2_,2) is complex, the topology of the graphs *DB*(*Reads*_1_,3) and *DB*(*Reads*_2_,3) depends on the read-set. *DB*(*Reads*_1_,3) consists of two connected components (top) while *DB*(*Reads*_2_,3) is a single-component graph since it has an extra edge (labeled **AGGC)** introduced by the additional read **AAGGCT**. **(C) Limitation of the de Bruijn graph approach to genome assembly.** Reckless resolution of unpaired complex vertices might disconnect the genome traversal. For a read-set *Reads*={**AAGGAG**, **AAGGCT**, **TTGG**}, since the read-path of **TTGG** ends inside the complex vertex **GG**, the vertex **TGG** (with red outline) represents a dead-end. **(D) Multiplex de Bruijn graph transformation.** The multiplex de Bruijn graph for the same read-set avoids reckless resolution of complex vertices by freezing unpaired complex vertices.

#### Path-graph

We say that a path *traverses* a vertex *w* in a graph if it both enters and exits this vertex. Given a path-set *Paths* in a graph, we denote the set of all paths containing an edge (*v,w*) as *Paths*(*v,w*) and the set of all paths traversing a vertex *w* as *Paths*(*w*). We define the set *Transitions*(*Paths*) as the set of all pairs of consecutive edges in all paths from *Paths*. A *path-graph G*(*Paths*) of a path-set *Paths* is defined as the transition-graph *G*(*Transitions*(*Paths*)).

Let *Paths* be the set of all read-paths in the compressed de Bruijn graph *G=DB*(*Reads*,*k*). A straightforward approach to constructing the graph *G*(*Paths*), that recomputes labels from scratch at each iteration, nearly doubles the path lengths at each iteration and thus faces the time/memory bottleneck. multiplexDBG avoids this time/memory bottleneck by modifying rather than recomputing the edge labels from scratch as described in Supplementary Note 10.

#### Multiplex de Bruijn graph transformation

Given a path-set *Paths* in a graph *G*, we call edges (*v,w*) and (*w,u*) in *G paired* if the transition-set *Transitions*(*Paths*) contain this pair of edges. A vertex *w* in *G* is *paired* if each edge incident to *w* is paired with at least one other edge incident to *w*, and *unpaired*, otherwise.

The important property of *DB*(*Genome,k*) is that there exists a genome traversal of this graph. Given a genome traversal of the graph *DB*(*Reads,k*), we want to preserve it in *DB*(*Reads,k*+1) after the graph transformation. However, it is not necessarily the case since the transformation of *DB*(*Reads,k*) into *DB*(*Reads,k*+1) may create dead-ends (each unpaired vertex in *DB*(*Reads,k*) results in a dead-end in *DB*(*Reads,k*+1)), thus “losing” the genome traversal that existed in *DB*(*Reads,k*) (Figure 5.C). Below we describe a single iteration of the algorithm for transforming *DB*(*Reads,k*) into the multiplex de Bruijn graph *MDB*(*Reads,k*) that avoids creating dead-ends whenever possible by introducing vertices of sizes (*k*+1) in this graph.

multiplexDBG transforms each paired vertex of *DB*(*Reads,k*) using the graph transformation algorithm and “freezes” each unpaired vertex by preserving its *k-*mer label and the local topology. It also freezes some vertices adjacent to the already frozen vertices even if these vertices are simple. Specifically, if a frozen vertex *u* is connected with a non-frozen vertex *v* by an edge of length *VertexSize(v)* + 1, multiplexDBG freezes *v* (Figure 5.D). The motivation for freezing *v* is that, if we did not freeze it, we would need to remove the edge connecting *u* and *v* in *MDB*(*Reads,k*), disrupting the topology of the graph. multiplexDBG continues the graph transformations for all paired vertices (while freezing unpaired vertices) with gradually increasing *k*-mer sizes from *k* to *K^+^*, resulting in the multiplex de Bruijn graph *MDB*(*Reads,k*) with *k*-mer varying in sizes from *k* to *K^+^*. Supplementary Note 11 illustrates that multiplex transformations may be overly-optimistic (by transforming vertices that should have been frozen) and overly-pessimistic (by freezing vertices that should have been transformed).

### Step 10: Expanding HPC contigs

Although LJA enables an accurate LJA assembly of HPC reads into HPC contigs (edge-labels in the multiplex de Bruijn graph), these contigs have to be expanded (de-collapsed) using information about homopolymer runs in the original reads. Supplementary Note 12 describes how LJApolish expands HPC contigs.

## Conclusions

The development of assembly algorithms for short reads (e.g., reads generated by Sanger and Illumina technologies) started from applications of the overlap/string graph approach. Even though this approach becomes slow and error-prone with respect to detecting overlaps in the highly-repetitive regions, the alternative de Bruijn graph approach (Idury and Waterman, 1995, Pevzner et al., 2001) was often viewed as a theoretical concept rather than a practical method.

Even after it turned into the most popular method for assembling short reads, the development of algorithms for assembling long error-prone reads (e.g., reads generated by Pacific Biosciences and Oxford Nanopore technologies) again started from the overlap/string graph approach (Koren et al., 2012, Chin et al., 2013, 2016) since the de Bruijn graph approach was viewed as inapplicable to error-prone reads due to the “error myth” (Roberts et al., 2013), Indeed, since long *k*-mers from the genome typically do not even occur in error-prone reads, it seemed unlikely that the de Bruijn graph approach may assemble such reads. However, the development of the Flye (Kolmogorov et al., 2019) and wtdbg2 (Ruan and Li, 2020) long-read assemblers demonstrated once again that the de Bruijn graph-based assemblers result in accurate and order(s) of magnitude faster algorithms than the overlap/string graph approach.

Since the de Bruijn graph approach was initially designed for assembling accurate reads, it would seem natural to use it for assembling long and accurate reads. However, the history repeated itself and the first HiFi assemblers again relied on the overlap/string graph approach (Nurk et al., 2020, Cheng et al., 2021). We described an alternative de Bruijn graph approach for assembling HiFi reads, illustrating that the “contest” between the de Bruijn graph approach and the overlap/string graph approach continues. Benchmarking on the T2T dataset demonstrated that LJA improves on the state-of-the-art HiFi assemblers with respect to both contiguity and accuracy. Although it is unclear how to conduct rigorous benchmarking without validated complete reference genomes, LJA results on the HG002 dataset illustrate that it generates highly contiguous phased assemblies. While this paper focuses on phased assemblies, it has immediate implications for the downstream applications since phased assemblies represent a stepping stone for both consensus and haplotype-resolved assemblies. For example, a straightforward bulge-collapsing and tip removal in the phased LJA assembly of the HG002 read-set results in a contiguous consensus assembly with N50=54 Mb. We are now developing the consensusLJA tool that will further increase the contiguity of this assembly, the diploidLJA tools for haplotype-resolved assembly, and the nanoLJA tool for constructing the de Bruijn graph from the latest generation of Oxford Nanopore reads with improved accuracy.

## Code Availability

The LJA code is available at https://github.com/AntonBankevich/LJA.

## Acknowledgments

Anton Bankevich, Andrey Bzikadze, and Pavel Pevzner were supported by the NSF EAGER award 2032783. Dmitry Antipov was supported by Saint Petersburg State University, Russia (grant ID PURE 73023672).

## Supplementary Notes

### Supplementary Note 1: De Bruijn graphs

Given a string-set *Genome* (each string in this set is either linear or cyclic and is referred to as a *chromosome*) and an integer *k*, the *de Bruijn multigraph* is defined as follows. Each *k*-mer from *Genome* corresponds to a vertex in the de Bruijn multigraph (identical *k*-mers correspond to the same vertex) and each (*k*+1)-mer corresponds to an edge connecting its *k*-prefix with its *k*-suffix (identical (*k*+1)-mers form parallel *edges* in the multigraph). The *uncompressed de Bruijn graph UDB*(*Genome,k*) is obtained from the de Bruijn multigraph by substituting each set of *m* parallel edges with a single edge of *multiplicity m*. Given a path *P* formed by edges *e_1_,e_2_,…,e_n_* in *UDB*(*Genome,k*), its *path-label label*(*P*) is defined as *label*(*e_1_*)\**lastSymbol*(*e_2_*)*…\**lastSymbol*(*e_n_*), where *lastSymbol*(*e*) stands for the last symbol of *label*(*e*) and *x*y* stands for the concatenate of strings *x* and *y*. We say that a path *P spells label*(*P*). A string-set *Genome* corresponds to a path-set that traverses each edge in the de Bruijn multigraph exactly once (and each edge in *UDB*(*Genome,k*) at least once) and spells *Genome*. We refer to this path-set as the *genome traversal*.

Given a read-set *Reads*, the uncompressed de Bruijn graph *UDB*(*Reads,k*) is constructed in the same way as *UDB*(*Genome,k*) by assuming that each read is a mini-chromosome. For a *k*-complete read-set, *UDB*(*Reads,k*)*=UDB*(*Genome,k*). In the case of error-prone reads, the goal is to construct the graph *UDB*(*Reads,k*) and modify it to approximate the graph *UDB*(*Genome,k*) as closely as possible.

Given a non-branching path *P* between junctions *v* and *w* in a graph, its *compression* results in substituting this path with a single edge (*v,w*) labeled by *label*(*P*). Compressing all non-branching paths in *UDB*(*Genome*,*k*) results in the *compressed de Bruijn graph DB*(*Genome*,*k*).

For an edge (*v*,*w*), an *edge*-*subpartition* substitutes this edge with two edges by “adding” an intermediate vertex *u* on this edge, i.e., deleting the edge (*v,w*) and adding a new vertex *u* to the graph along with edges (*v,u*) and (*u,w*). A *subpartition* of a graph is defined as a result of a series of subpartitions.

### Supplementary Note 2: Information about datasets

Supplementary Table S1 describes T2T, HG002, MOUSE, MAIZE, and FLY read-sets. We also analyze the T2TX dataset, a subset of the T2T dataset that contains all reads originating from chromosome X.

**Supplementary Table S1.** Information about HiFi read-sets used for LJA benchmarking. The total length of the homopolymer-collapsed *T2TGenome* is 2,133,004,165. The T2TX read-set was generated by mapping the T2T read-set to *T2TGenome* with Winnowmap (Jain, 2020) and selecting reads mapped to chrX. In rare cases, when a read maps to multiple nearly identical instances of a repeat, Winnowmap outputs both *primary* and *secondary* read alignments. Although using primary alignments works well for a vast majority of regions in the human genome, primary alignments may incorrectly map some reads in the most repetitive regions such as centromeres. These incorrectly mapped reads may result in low coverage of some repeat instances, thus negatively affecting the generation of datasets containing error-free reads. We thus used both primary and secondary alignments in such regions. The T2TErrorFree dataset was derived by mapping reads from the T2T dataset to *T2TGenome* and substituting each of 5,567,034 mapped reads by the genomic segment it spans.

| dataset | #reads | median read length (kbp) | coverage | genome length (Gb) | origin |
| --- | --- | --- | --- | --- | --- |
| T2T | 5,567,158 | 17.2 | 32 | 3.1 | SRX7897688<br>SRX7897688<br>SRX7897688<br>SRX7897688 |
| T2TX | 272,732 | 17.2 | 32 | 0.15 | T2T |
| HG002 | 7,384,171 | 15.0 | 36 | 3.1 | SRP227894 |
| FLY | 2,390,566 | 10.2 | 120 | 0.2 | SRX8020156 |
| MOUSE | 4,060,680 | 16.4 | 26 | 2.6 | SRR11606870 |
| MAIZE | 3,089,410 | 15.6 | 20 | 2.4 | SRR11606869 |

### Supplementary Note 3: Benchmarking individual modules of the LJA pipeline

#### Benchmarking jumboDBG

Supplementary Table S2 presents information about the running time and memory footprint of jumboDBG. Even though the de Bruijn graphs *DB*(*T2TGenome,k*) of the human genome greatly vary in complexity when the *k*-mer size varies, jumboDBG constructs each of these graphs in ≅40 min with ≅35 Gb memory footprint for all *k*-mer sizes (Supplementary Figure S1).

**Supplementary Figure S1.**
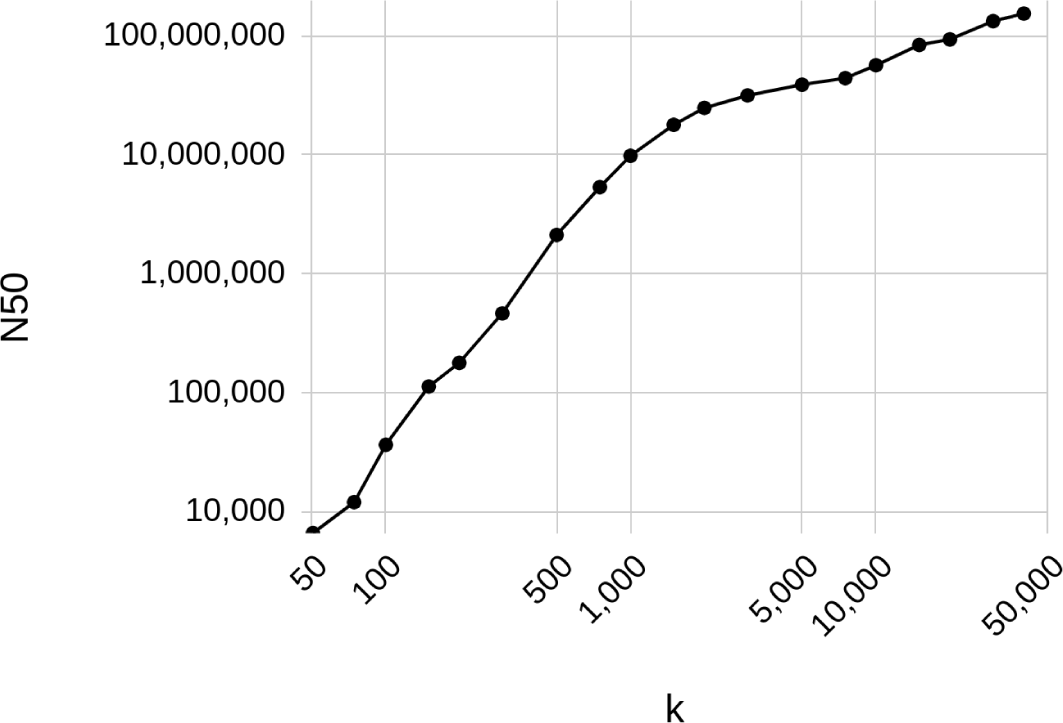
Information about the graph *DB*(*T2TGenome, k*) for *k* varying from 50 to 40000. The edge-labels in the compressed de Bruijn graph define a set of strings (contigs). The figure presents the N50 metric of the contigs that represent edge-labels in *DB*(*T2TGenome, k*). At *k*=20000, the graph has only 401 edges (N50=93 Mb) with 5 out of 23 chromosomes completely assembled. At *k*=40000, the graph has only 116 edges (N50=154 Mb) with 16 out of 23 chromosomes completely assembled. Note that N50=154Mb for contigs formed by the entire human chromosomes.

**Supplementary Table S2.** The running time and memory footprint of jumboDBG in the compressed de Bruijn graphs for the T2T (top) and T2TErrorFree (bottom) read-sets. The T2TErrorFree read-set was derived by mapping reads from the T2T dataset to *T2TGenome* and substituting each mapped read by the genomic segment it spans. The table provides information about the number of vertices and edges in the graph, the number of distinct read-paths (excluding single-edge read-paths), their median length (in the number of edges), and the number of complex vertices in the graph. The running time and memory footprint for the T2TErrorFree read-set hardly changes with an increase in the *k*-mer size, suggesting that they mainly depend on the size of the compressed de Bruijn rather than the *k*-mer size. All information is provided for HPC reads. All tools were benchmarked on a computational node with two Intel Xeon 8164 CPUs, with 26 cores each and 1.5 TB of RAM. All runs were done in 32 threads.

| T2T |  |  |  |  |  |  |  |  |
| --- | --- | --- | --- | --- | --- | --- | --- | --- |
| $k$ | time (h) | memory (Gb) | #vertices | #edges | total edge length (Gbp) | #read-paths | median read-path length | #complex vertices |
| 251 | 2.4 | 52 | 29806686 | 43750620 | 10 | 5566455 | 159 | 208737 |
| 501 | 2.7 | 54 | 23171326 | 33230906 | 15 | 5566321 | 111 | 95702 |
| 1001 | 3.2 | 65 | 17032673 | 23357700 | 18 | 5559705 | 69 | 36041 |
| 2001 | 3.6 | 81 | 12025225 | 14947471 | 23 | 5422657 | 35 | 11493 |
| 3001 | 4.2 | 90 | 9927841 | 11160720 | 25 | 5157197 | 22 | 5308 |
| 5001 | 3.9 | 93 | 8152006 | 7560390 | 27 | 4564678 | 10 | 1581 |
| T2ErrorFree |  |  |  |  |  |  |  |  |
| $k$ | time (h) | memory (Gb) | #vertices | #edges | total edge length (Gbp) | #read-paths | median read-path length | #complex vertices |
| 251 | 0.6 | 32 | 311042 | 472380 | 2.1 | 874719 | 55 | 7432 |
| 501 | 0.6 | 33 | 143011 | 214517 | 2.1 | 590730 | 30 | 1230 |
| 1001 | 0.7 | 33 | 64371 | 95716 | 2.1 | 456338 | 13 | 220 |
| 2001 | 0.7 | 34 | 21724 | 31862 | 2.0 | 307824 | 5 | 16 |
| 3001 | 0.6 | 35 | 10035 | 14723 | 2.0 | 205879 | 4 | 8 |
| 5001 | 0.6 | 36 | 3530 | 4956 | 2.0 | 88460 | 3 | 2 |

#### Benchmarking mowerDBG

Error-correction of reads, introduced in Pevzner et al., 2001, has become ubiquitous in both short-read and long-read assemblers (Chaisson et al., 2008, Kelly et al., 2010, Medvedev et al., 2011, Bankevich et al., 2012, Nikolenko et al., 2013, Lima et al., 2020). However, error-correction of long and accurate reads remains a poorly explored topic — hifiasm (Cheng et al., 2020) is currently the only error-correcting tool for HiFi reads. The hifiasm error-correction of the T2T dataset resulted in only 5.3 errors per a megabase of the total read-length and decreased the percentage of erroneous reads to 3.2%. However even with this seemingly very small error rate, the de Bruijn graphs on error-corrected reads become inadequate when the *k*-mer size is large.

mowerDBG significantly reduces the error rate in reads as compared to hifiasm (from 5.3 to 2.4 errors per megabase) and decreases the percentage of error-prone reads to 2.6% (after “breaking” a small number of reads as described in Supplementary Note 8). Moreover, most of the remaining errors reflect the pseudo-heterozygosity in the CHM13 cell line (Nurk et al., 2021) rather than real errors. This improved error correction in mowerDBG is critical for enabling the de Bruijn graph approach.

#### Benchmarking multiplexDBG on human chromosome X

We benchmarked multiplexDBG using the T2TX read-set, the subsets of the T2T dataset that contains all reads originating from human chromosome X. We also analyzed the T2TXErrorFree dataset obtained by substituting each read in T2TX by the segment of the genome it spans. We constructed the graph *DB*(*T2TXErrorFree,k*) of the read-set *T2TXErrorFree* and the graph *DB*(*chrX,k*) of chromosome X for *k*-mer sizes varying from 1001 to 40001. Additionally, we applied multiplexDBG to the graph *DB*(*T2TXErrorFree,k*), resulting in the multiplex de Bruijn graph *MDB*(*T2TXErrorFree,k*).

We define the size of a graph *G* as the number of its edges and denote it as |*G*|. Supplementary Figure S2 illustrates that multiplexDBG significantly reduces the size of the de Bruijn graph (and increases the contiguity of the assembly) when it transforms it into the multiplex de Bruijn graph, Both *DB*(*T2TXErrorFree,k*) and *MDB*(*T2TXErrorFree,k*) represent approximations of *DBG*(*chrX, k),* albeit with different accuracy. We note that, in addition to all *k*-mers from *T2TXErrorFree* read-set, *MDB*(*T2TErrorFree,k*) uses virtual reads to recover some *k*-mers from chromosome X missing in this read-set. Supplementary Figure S2 illustrates that |*DB(chrX,k)*| *≤* |*MDB(T2TXErrorFree, k)*| *≤* |*DB(T2TXErrorFree,k)*| until the *k*-mer size exceeds *k*≅40001, a point when the graph *DB(T2TXErrorFree,k)* rapidly disintegrates since there are very few reads longer than 40001.

**Supplementary Figure 2.**
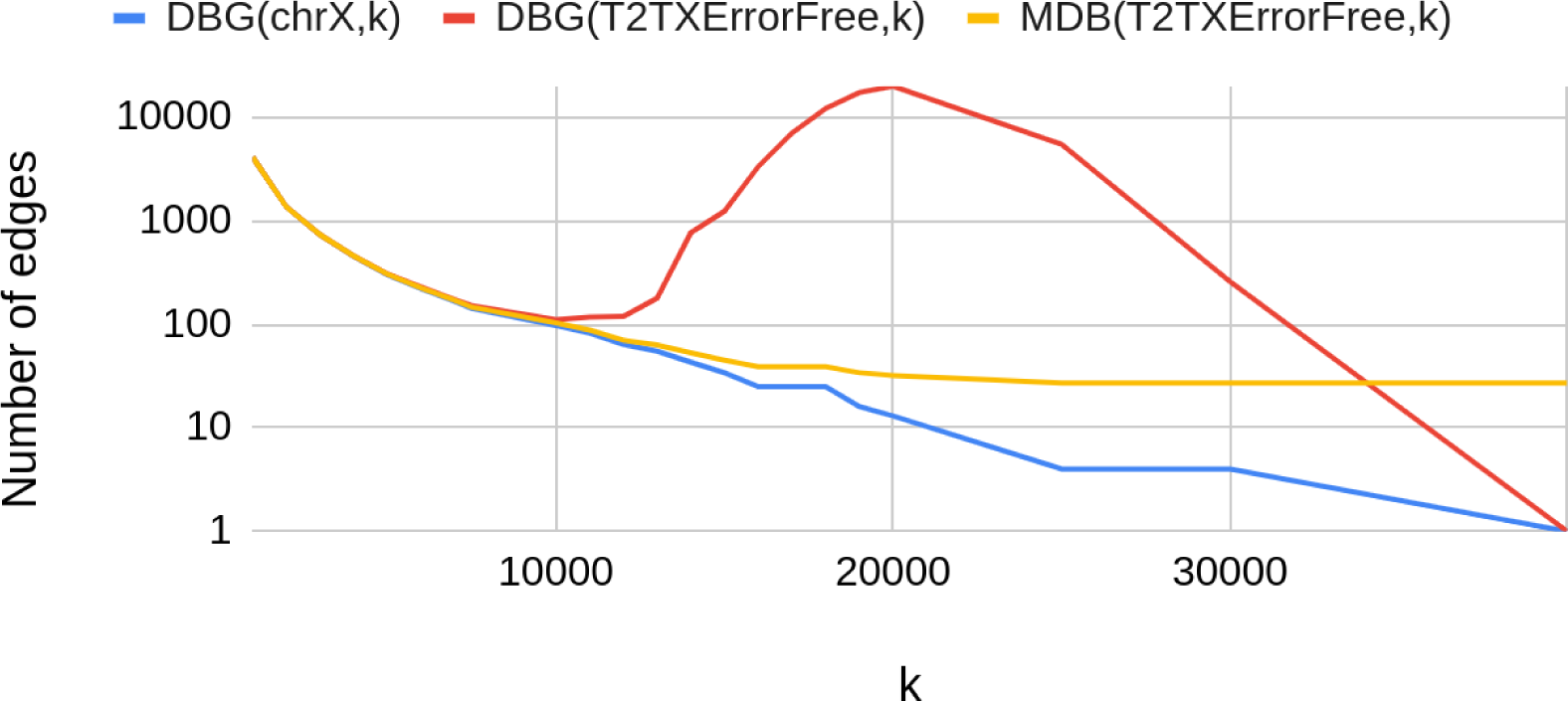
Comparison of the sizes of the compressed de Bruijn graph and the multiplex de Bruijn graph. The plots shows the sizes on the de Bruijn graph *DB*(*chrX,k*) (blue), *DB(T2TXErrorFree,k)* (red), and *MDB(T2TXErrorFree,k)* (yellow) of the same read-set for *k* ranging from 1001 to 40001. All three graphs are identical for *k* < 4001 since the read-set *T2TXErrorFree* contains all genomic 4001-mers from chromosome X. Since more and more genomic *k*-mers are missing from the *T2TXErrorFree* read-set with increasing *k*-mer size, the red and blue curves diverge for *k* > 4001, reflecting the increasing size of *DB*(*T2TXErrorFree,k).* However, the graph *MDB*(*T2TErrorFree,k*) successfully recovers the missing *k*-mers and continues to perfectly match *DB*(*chrX,k*) for *k <* 7501. When the *k*-mer size further increases, information in reads becomes insufficient for resolving some long repeats and the size of *MDB*(*T2TXErrorFree,k)* starts exceeding the size of *DB*(*chrX,k*), resulting in the diverged yellow and blue curves. Since the T2T dataset has very few reads longer than *k=*25001, the size of *DB*(*T2TXErrorFree,k*) rapidly drops and the size of *MDB(T2TXErrorFree,k)* stabilizes when the *k*-mer size exceeds *≅*25001.

All three graphs *DB(T2TXErrorFree,k)*, *MDB(T2TXErrorFree,k),* and *DB(chrX,k),* are identical when the *k*-mer size does not exceed 4501. For larger *k-*mers, *DB(T2TXErrorFree,k)* starts to suffer from breaks in *k*-mer coverage that fragment edges of the de Bruijn graphs. In contrast, the multiplex de Bruijn graph recovers all missing *k*-mers and provides a perfect representation of *DB(chrX,k)* until *k*≅7501 when the gaps in coverage start to occasionally fall into complex repeats. Still, *MDB(T2TXErrorFree,k)* represents an excellent approximation of *DB*(*ChrX,k*) until *k*=15001. At *k*≅25001, since the number of reads longer than 25001 in the T2T dataset rapidly decreases, the size of *DB*(*T2TXErrorFree,k)* rapidly drops. In contrast, the size of *MDB*(*T2TXErrorFree,k*) stabilizes starting at *k*≅25001 since it preserves the structure of the graph constructed from smaller *k*-mers and has no additional information for further graph transformations.

In total, multiplexDBG reduced the size of the compressed de Bruijn graph *DB*(T2T, 5001) from 6516 to 1440 edges. Below we describe additional benchmarking of multiplexDBG on human centromeres.

#### Benchmarking multiplexDBG on human centromeres

For a centromere on chromosome N, we consider a subset of the T2T dataset that contains all reads originating from this centromere (referred to as T2TcenN). We denote the error-free version of this dataset as T2TcenNErrorFree. Below we analyze human centromeres X (6527 reads) and 6 (11409 reads), each of length ≅3 Mb.

Extremely repetitive cen6 is one of the most repetitive regions of the human genome that was first assembled using ultralong Oxford Nanopore reads (Bzikadze and Pevzner, 2020). Assembling cen6 using shorter HiFi reads is challenging since it contains long nearly identical repeats. Below we illustrate how the multiplex de Bruijn graph approach results in a nearly complete cenX assembly from the real T2TcenX read-set. In contrast, it is impossible to assemble cen6 even from the error-free T2Tcen6ErrorFree read-set, illustrating that ultralong reads are needed to generate telomere-to-telomere assemblies.

The graph *DB*(*T2TcenXErrorFree,*5001) contains 34 vertices and 49 edges. After increasing the k-mer size from 5001 to 40001, multiplexDBG transformed this graph into the multiplex de Bruijn graph *MDB*(*T2TcenXErrorFree,*5001) with only three edges (contigs). Assembling centromere X from real reads resulted in the same graph *MDB*(*T2TcenX,*5001).

In contrast, the graph *DB*(*T2TcenXErrorFree,*40001) cannot be constructed, because all reads in T2TcenXErrorFree are shorter than 40001 bp. We emphasize that the multiplex de Bruijn graph utilizes virtual reads (Supplementary Note 4) that are often longer than real HiFi reads, explaining why it was important to increase the *k^+^*-mer size beyond the length of all reads in the T2TcenXErrorFree read-set.

The more complex graph *DB*(*T2Tcen6ErrorFree,*5001) of centromere 6 contains 152 vertices and 226 edges (the read-set T2Tcen6ErrorFree contains no missing genomic 5001-mers). multiplexDBG transformed this graph into a small multiplex de Bruijn graph *MDB*(T2Tcen6ErrorFree,5001) with only 10 vertices and 15 edges in ≅2 minutes (Supplementary Figure S3). The large reduction in complexity of *MDB*(*T2Tcen6ErrorFree,*5001) as compared to *DB*(T2Tcen6ErrorFree,5001) illustrates the value of multiplex de Bruijn graphs for the follow up repeat resolution using ultralong reads.

**Supplementary Figure S3.**
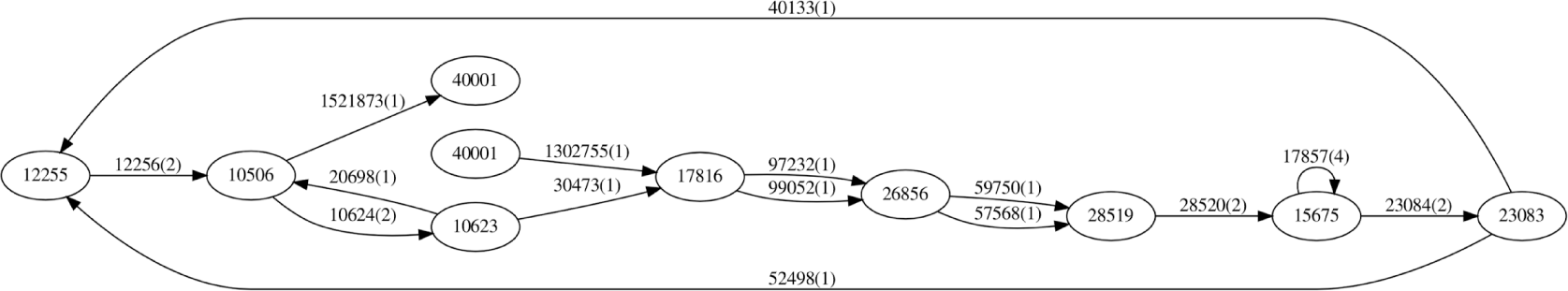
The multiplex de Bruijn graph *MDB*(*T2Tcen6ErrorFree*,5001). The length and multiplicity of each edge is shown next to this edge.

### Supplementary Note 4: Virtual reads

A path (*v_1_,…v_i_,…,v_n_*) in a graph is called *out-unambiguous* (*in-unambiguous*) if the outdegrees (indegrees) of all vertices in this path except the first and the last one are equal to 1. A path (*v_1_,…v_i_,v_i+1_,…,v_n_*) is called *unambiguous* if there is an edge (*v_i_,v_i+1_*) in this path such that (*v_i_,…,v_n_*) is an *out-unambiguous* path and (*v_1_,…v_i+1_*) is an *in-unambiguous* path (Supplementary Figure S4). We refer to unambiguous paths in the compressed de Bruijn graph as *virtual reads*. Note that in the case when *Genome* is formed by circular chromosomes, all virtual reads in the compressed de Bruijn graph *DB*(*Genome,k*) represent substrings of *Genome* and thus can be safely added to any read-set. In the case of linear chromosomes, we assume that the *k*-prefix and *k*-suffix of each chromosome correspond to dead-ends in the graph *DB*(*Reads, k*). Virtual reads are important for multiplex de Bruijn graph construction since multiplexDBG complements the error-corrected read-set *Reads\** by adding all virtual reads derived from the de Bruijn graph *DB*(*Reads*,K*).

**Supplementary Figure S4.**
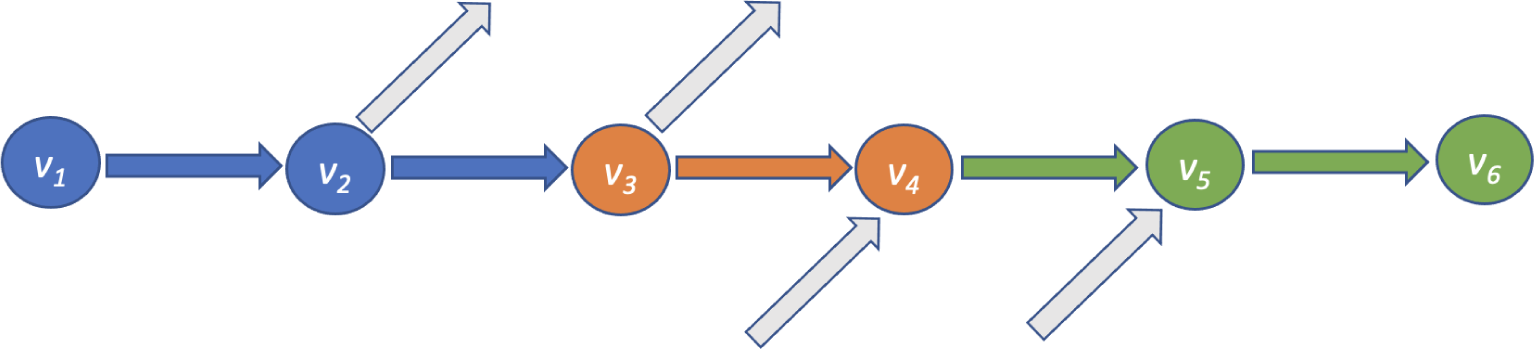
An unambiguous path. Since vertices *v_2_,v_3_* have indegrees 1 and vertices *v_4_,v_5_* have outdegrees 1, the path (*v_1_,v_2_,v_3_*) is in-unambiguous and the path (*v_3_,v_4_,v_5_*) is out-unambiguous. Thus, the joined path (*v_1_,v_2_,v_3_,v_4_,v_5,_ v_6_*) is unambiguous.

### Supplementary Note 5: Analyzing variations in genome coverage by reads

#### Coverage variations in the T2T dataset

Below we consider the error-corrected read-set T2T* (after wto rounds of error-correction). We define the *normalized coverage* of a *k*-mer (edge) in the de Bruijn graph as its coverage by reads divided by the average coverage across all *k*-mers in the genome. We define *cov_k_*(*x*) (*edge-cov_k_*(*x*)) as the fraction of *k*-mers (edges) with normalized coverage below *x* among all *k*-mers (edges) in the de Bruijn graph. The normalized coverage of the vast majority of *k*-mers and edges in the graph *DB*(*T2T*,*501) falls in the interval between 0.5 and 1.5, e.g., *cov_501_*(0.5)=0.006, *cov_501_*(1.5)=0.98, *edge-cov_501_*(0.5)=0.01, and *edge-cov_501_*(1.5)=0.83 (Supplementary Figure S5).

**Supplementary Figure S5.**
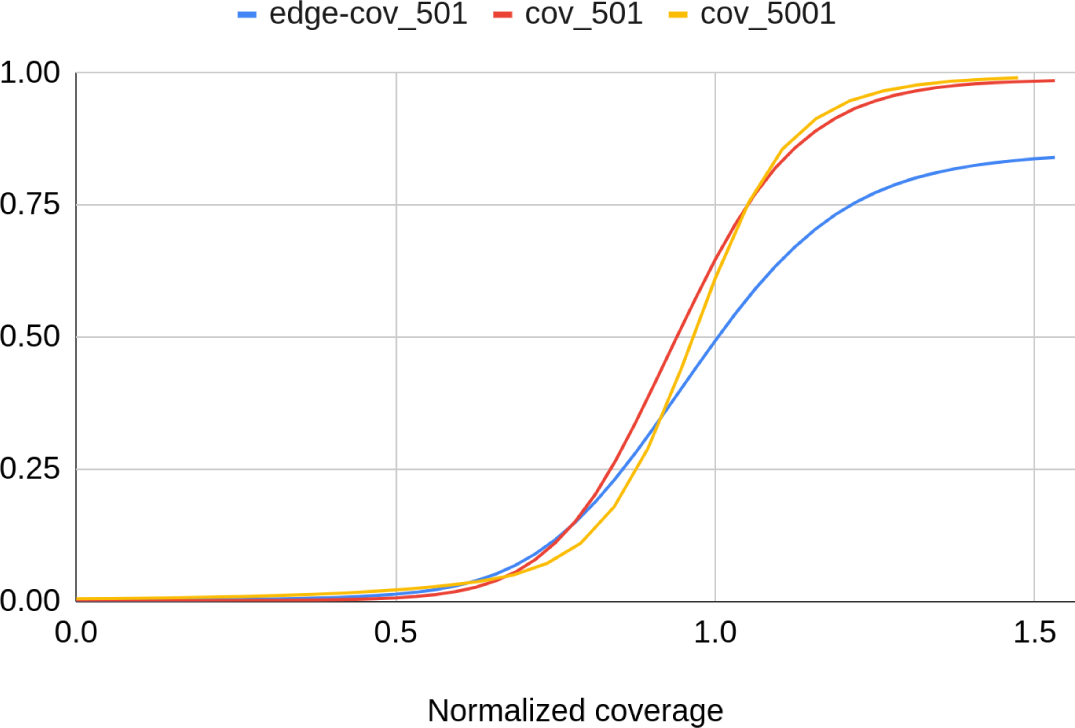
Functions *cov_501_*(*x*) and *cov_5001_*(*x*), as well as the function *edge-cov_501_*(*x*), for the error-corrected read-set T2T*. The average coverage across all 501-mers (5001-mers) in *T2TGenome* is 32 (19).

To analyze how the coverage of an edge correlates with its multiplicity, we mapped each edge *e* in the de Bruijn graph *DB*(*T2T*,k*) of reads to a most similar edge *e’* in the de Bruijn graph *DB*(*T2TGenome,k*) of genome and computed the ratio of the normalized coverage of *e* and the multiplicity of *e’*. Supplementary Figure S6 shows the distribution of this value over all edges of the graph *DB*(*T2T*,*501) and illustrates that the coverage can be used as a rough estimate of the multiplicity for most edges (the ratio falls in the interval from 0.9 to 1.1 for the vast majority of edges).

**Supplementary Figure S6.**
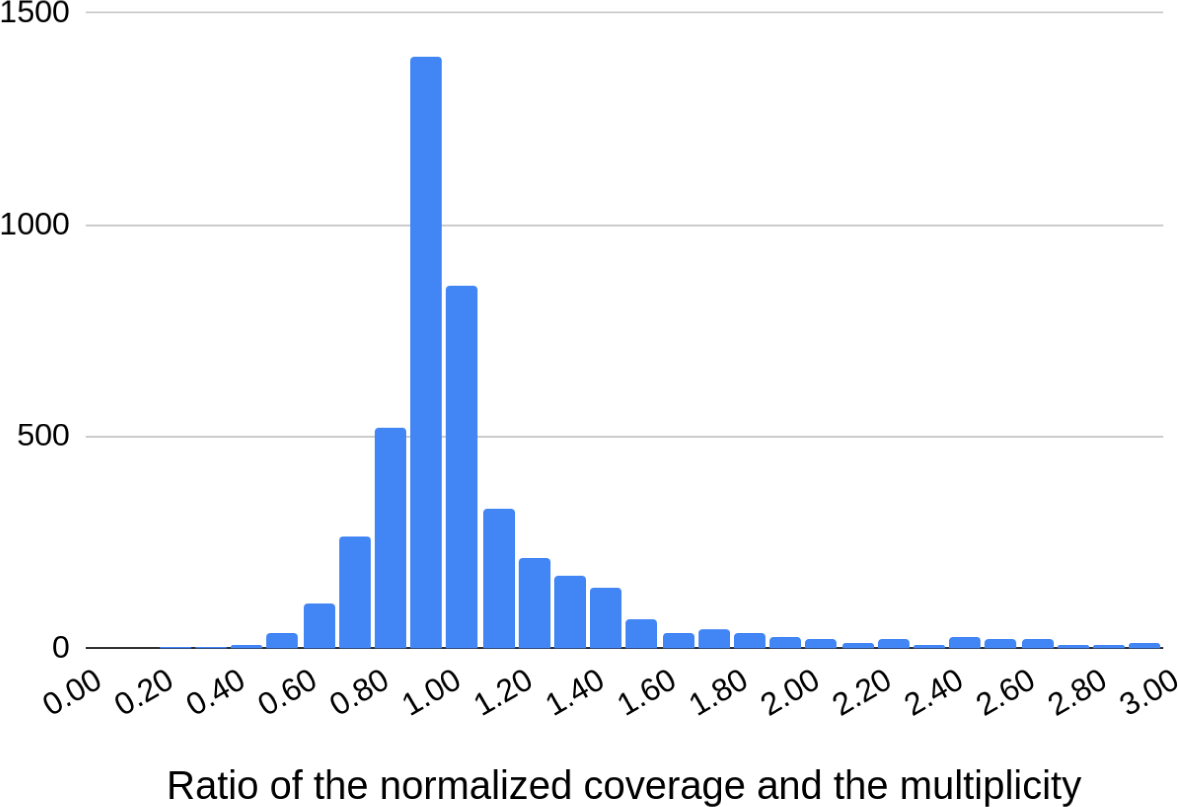
Comparing the normalized coverage of edges in *DB*(*T2T*,501*) with the multiplicities of the corresponding edges in *DB*(*T2TGenome,501*). The histogram is generated for edges of length at least 5000 bp as the coverage estimates for shorter edges are less reliable. Each bin of size 0.1 shows the number of edges with the given ratio of the normalized coverage and the multiplicity.

#### Coverage variation in the FLY dataset

Supplementary Figure S7 presents information about the coverage variation in the FLY dataset.

**Supplementary Figure S7.**
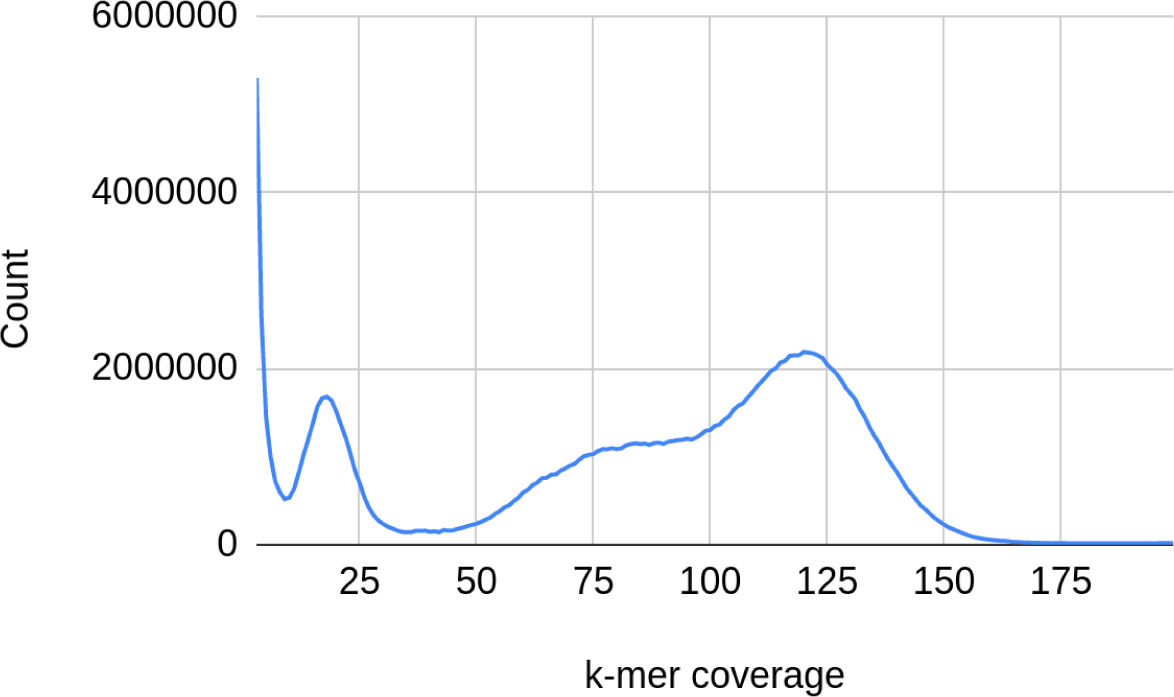
Distribution of coverage of 501-mers in the FLY read-set. Although the coverage of 501-mers by the FLY read-set is ≅120x, there is a large number of *k*-mers with coverage ≅60x, pointing to some diverged maternal and paternal alleles in this inbred genome. A peak at coverage ≅20x likely represents 501-mers from a bacterial contaminant.

### Supplementary Note 6: LJA parameters

#### jumboDBG parameters

Below we describe jumboDBG parameters.

#### Bloom filter

The parameters *BloomSize* and *BloomNumber* define the tradeoff between the memory/query time of the Bloom filter and its false positive rate. For example, increasing the number of hash functions decreases the false positive rate but increases the memory footprint and the query time of the Bloom filter. jumboDBG selects a rather small default value *BloomNumber*=5 and sets the parameters *BloomSize* in such a way that the false positive rate of the Bloom filter does not exceed the threshold *FPrate* with the default value 10^-4^.

Given the expected number *N* of of different elements to be stored in a Bloom filter, its false positive rate is estimated as (1-*e^-BloomNumber* N / BloomSize^*)*^BloomNumber^* (Mitzenmacher and Upfal, 2005). When *BloomNumber* is fixed, the false positive rate depends only on the ratio *N/BloomSize*. Thus, *BloomSize* for constructing *DB*(*Genome,k*) should be selected proportional to the number of different (*k*+1)-mers in *Genome* (*N*) in such a way that the false positive rate of the Bloom filter does not exceed the *FPrate* threshold. Similarly, *BloomSize* for constructing *DB*(*Reads,k*) should be selected proportional to the number of different (*k*+1)-mers in *Reads.* Below we describe how jumboDB estimates the number of different (*k*+1)-mers in *Genome* and *Reads*.

Given a string-set *S*, we define *length_k_*(*S*) as the total number of (*k*+1)-mers in this string-set (the number of (*k*+1)-mers in an *n*-nucleotide cyclic (linear) string is *n* (*n-k*)). jumboDB sets the parameter Since the number of different (*k*+1)-mers in *Genome* is unknown, jumboDBG uses *length_k_*(*Genome*) as a proxy for this number during construction of *DB*(*Genome,k*). Since the number of different (*k*+1)-mers in *Reads* is also unknown, jumboDBG uses *length_k_*(*Disjointigs*) as a proxy for this number during construction of *DB*(*Reads,k*). As a tradeoff between the memory footprint and the false positive rate of the Bloom filter, we use 32 bits per a (*k*+1)-mer, resulting in the approximately ≅10^-4^ false-positive rate.

#### Rolling hash

jumboDBG uses a 128-bit polynomial rolling hash for storing (*k*+1)-mers in reads. Although hashing may lead to collisions when different (*k*+1)-mers result in the same hash function, a 128-bit rolling hash has a very low probability of collisions. Indeed, if a hash is viewed as a pseudo-random number generator on a set of 128-bit integers, the probability of a collision during the construction of the graph *DB*(*Genome*,*k*) is extremely low even for large genomes. Moreover, it remains small during the construction of the graph *DB*(*Reads*,*k*) - it is estimated as 10^-17^ in the case of the T2T read-set with approximately 10^11^ 501-mers. We performed many tests on (*k*+1)-mers from the T2T read-sets and detected no collisions.

#### Minimizers

Since jumboDBG discards all reads shorter than *width+k*, it selects parameter *width* in such a way that the vast majority of reads have length at least *width+k*. It thus selects *width* = 2000 for *k*=501 and *width* = 500 for *k*=5001.

### mowerDBG parameters

Since errors hardly ever trigger identical erroneous *k*-mers for a sufficiently large *k*-mer size (except for errors inside dinucleotide repeats), mowerDBG sets a rather small default threshold *minCoverage*=2.

Ideally, the *maxRepeatLength* threshold should be equal to the length of the longest perfect repeat in the genome. Since this length is unknown, mowerDBG sets a rather large threshold *maxRepeatLength*=40001. Increasing this threshold even further is unlikely to compromise the error-correction since the flows in the network algorithm in mowerDBG identifies unrepeated edges even if they are shorter than *maxRepeatLength*.

mowerDBG selects the *fraction* parameter (default value 0.01) to be slightly larger than the error rate in HiFi reads. It sets *coverageAmplifier=*1.5 because coverage of genomes by HiFi reads is rather uniform and rarely exceeds the median coverage by more than 50%.

### multiplexDBG parameters

multiplexDBG has a single parameter *K^+^* (default value = 40,001) to transform the compressed de Bruijn graph *DB*(*Reads,K*) into the multiplex de Bruijn graph *MDB*(*Reads,K*) with *k*-mer-sizes varying from *K* to *K^+^*.

### LJApolish parameters

To limit the memory footprint, LJApolish uses at most *maxReadNumber* reads for computing the median of multiplicities of homopolymers (default value 20). It uses parameters *bandwidth* (default value 10) and *bandwidthLarge* (default value 320) as limits for the bandwidth size in the banded alignment and uses the same parameter *long_dinucleotide* (default value 16) for collapsing dinucleotide repeats as in the initial read collapsing step of the LJA pipeline.

### Supplementary Note 7: Constructing a compact sparse de Bruijn graph

In contrast to the de Bruijn graph, with degrees bounded by the alphabet size, vertices of a sparse de Bruijn graph may have arbitrarily large degrees. Below we describe how jumboDBG transforms a sparse de Bruijn graph *SDB*(*Reads,Anchors*) into a compact sparse de Bruijn graph *SDB*(*Reads,Anchors\**) using an example of a vertex *w* that has one incoming edge *in* and two outgoing edges *out_1_* and *out_2_* (a similar approach is applicable to any vertex). It achieves this goal by “extending” the label of the edge *in* and “shortening” the labels of edges *out_1_* and *out_2_* (Supplementary Figure S8).

**Supplementary Figure S8.**
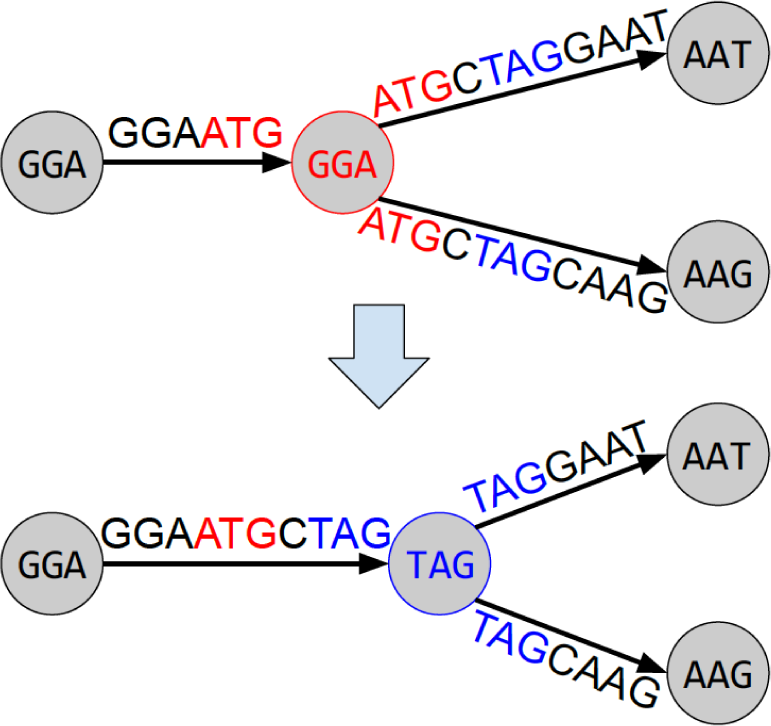
Transforming a sparse de Bruijn graph (top) into a compact sparse de Bruijn graph (bottom). Vertex *w* in the sparse de Bruijn graph (top) with red label ATG has one incoming edge *in* and two outgoing edges *out_1_* and *out_2._*. It is substituted by a vertex with blue label TAG after extending the edge-label of *in* and shortening the edge-labels of *out_1_* and *out_2_*. Since *prefix*(*out_1_,out_2_*)=ATGCTAG, edges *out_1_* and *out_2_* share their first five *k*-mers ATG, TGC, GCT, CTA, and TAG. Since TAG is the last *k*-mer of *prefix*(*out_1_,out_2_*), removing a non-junction vertex ATG and adding a new junction vertex TAG reduces the number of non-junction vertices in the sparse de Bruijn graph and eventually transforms it into the compact sparse de Bruijn graph.

Edges *out_1_* and *out_2_* share their first *k*-mer (that labels vertex *w*) and possibly their second, third, etc. *k*-mers. Let *prefix*(*out_1_,out_2_*) be the longest common prefix of these edges and *last*(*out_1_,out_2_*) be the last *k*-mer of this prefix. While the vertex *w* is not necessarily a junction, the edges *out_1_* and *out_2_* share a junction *last*(*out_1_,out_2_*). We thus extend the label of the edge *in* ending in *w* by concatenating it with the suffix of *prefix*(*out_1_,out_2_*) starting at position *k*, and shorten the labels of edges *out_1_* and *out_2_* starting in *w* by removing their prefixes of length |*prefix*(*out_1_,out_2_*)|-*k*. As the result of this operation, edges that previously started/ended at an anchor *w*, now start/end at a junction *last*(*out_1_,out_2_*). Applying the described procedure to all vertices of the graph *SDB*(*Reads,Anchors*) transforms it into a compact sparse de Bruijn graph *SDB*(*Reads,Anchors\**). Edge-labels in this graph form a compact disjointig-set.

### Supplementary Note 8: Error correction algorithm

An edge in the graph *DB*(*Reads,k*) is classified as *low-coverage* if its coverage is below a threshold *minCoverage*, and a *high-coverage*, otherwise. Low-coverage and high-coverage edges partition each read-path in the graph *DB*(*Reads,k*) into alternating *low-coverage* and *high-coverage subpaths*. The mowerDBG error-correction approach is motivated by the observation that nearly all high-coverage paths in *DB*(*Reads,k*) are formed by genomic (*k*+1)-mers, while nearly all low-coverage paths are caused by errors in reads.

Next subsection analyzes low-coverage and high-coverage paths in the de Bruijn graphs. Afterward, we describe how mowerDBG corrects reads by performing four steps: (i) read-rerouting, (ii) bulge-collapsing, (iii) correcting dimers, and (iv) correcting pseudo-correct reads. We use the graph *DB*(*T2TX*,501) to illustrate how mowerDBG error-corrected reads after each of these steps.

#### Analyzing low-coverage and high-coverage paths in the de Bruijn graph

Supplementary Figure S9 illustrates that the vast majority of *genomic edges* in *DB*(*T2TX*,501) (edges that are visited by the genome traversal) are high-coverage edges. This graph has only 31 isolated edges that correspond to reads that do not share 501-mers with other reads. In contrast, the graph *DB*(*T2TX*,5001) has many more isolated edges (1454). However, after constructing the graph *DB*(*T2TX\**,5001) on the error-corrected read-set, there are only 398 reads forming isolated edges in this graph.

**Supplementary Figure S9.**
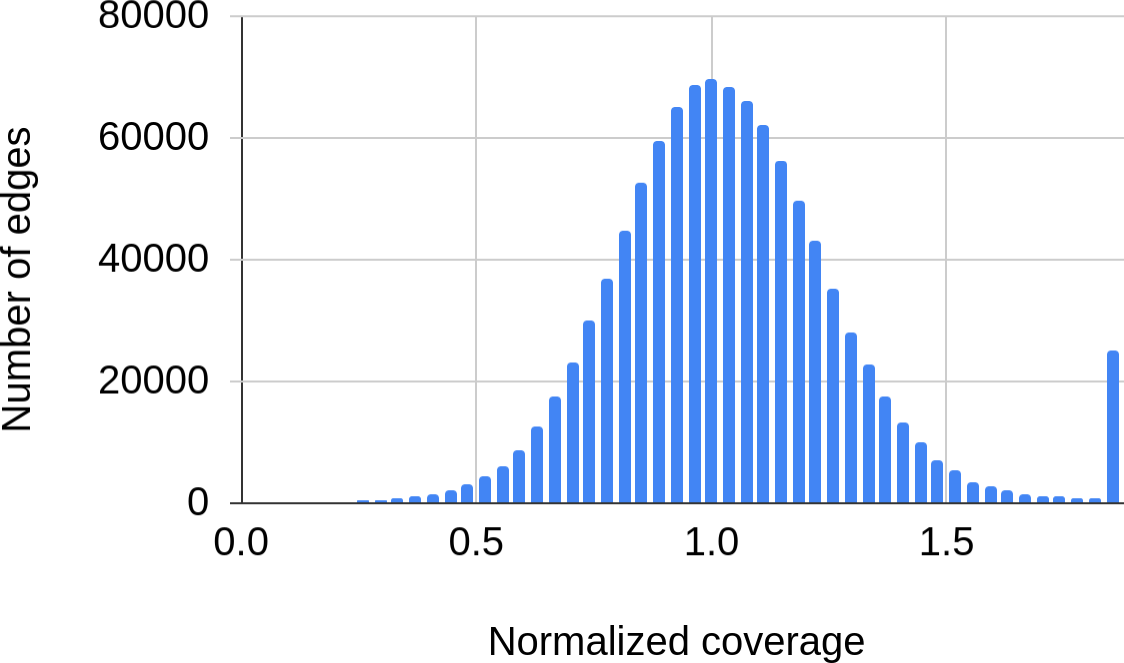
Histogram of the normalized coverage of genomic edges in the graph *DB*(*T2TX*,501). The normalized coverage of an edge is defined as the coverage of this edge divided by the median coverage of all edges in the graph. Each bar shows the number of edges in the bin of size 0.04. The histogram has a long tail formed by edges that are visited multiple times by the genome traversal and the bar at normalized coverage 2 shows the total number of edges with normalized coverage at least 2. The genome traversal of chromosome X is formed by 1,180,374 edges in *DB*(chrX,501) (only 56/71/112 of them have a small coverage 1/2/3 in *DB*(*T2TX*,501)). Since 6841 501-mers from the assembled chromosome X do not appear in reads and form 19 coverage gaps, the genome traversal in *DB*(*T2TX*,501) consists of 20 paths rather than a single path.

When a genome is known, one can align each read (corresponding to a read-path *P* in *DB*(*Reads*,*k*)) to the genome, find the genomic segment spanned by this read, and identify a path *P_Genome_* in *DB*(*Reads*,*k*) that spells this fragment. Although this procedure works for the vast majority of reads and typically results in a high-coverage path *P_Genome_*, this path often differs from the original path *P*. An edge in a read-path *P* is called *correct* if it is also an edge in *P_Genome_* (and the corresponding edges are aligned against each other in the read-genome alignment), and *incorrect*, otherwise. The correct and incorrect edges partition the read-path *P* into the correct and incorrect subpaths. Given an incorrect subpath *P\** of *P*, its *valid correction* is defined as substituting this subpath with a subpath of *P_Genome_* that *P\** is aligned to (all other corrections are classified as *invalid*). 98.2% of the incorrect subpaths in the T2TX dataset are low-coverage subpaths and 99.95% of low-coverage subpaths in the T2TX dataset are incorrect subpaths.

A subpath of a path *P* is called *external* if it contains the first or the last edge of *P*, and *internal*, otherwise. Supplementary Figure S10 illustrates that the vast majority of low-coverage internal subpaths in the graph *DB*(*Reads,k*) spell strings of length close to *k* and the vast majority of external subpaths are short.

**Supplementary Figure S10.**
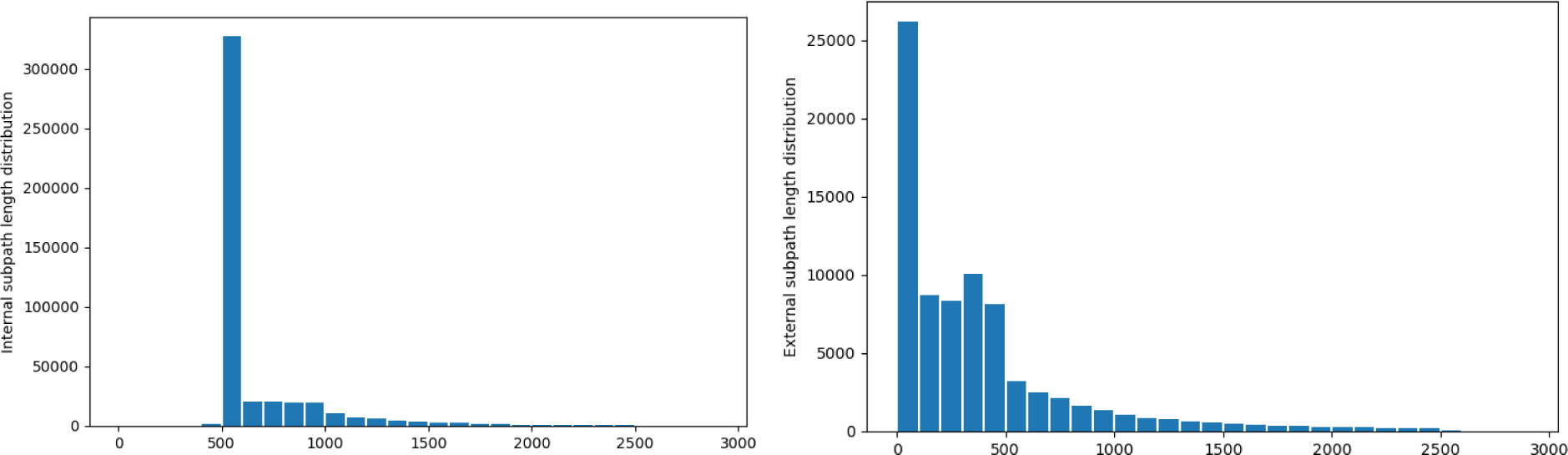
Distribution of lengths of strings spelled by all 455,548 internal (left) and 79,398 external (right) low-coverage subpaths of read-paths in the graph *DB*(T2TX,501).

#### Bypasses

We first describe how mowerDBG corrects internal low-coverage subpaths. Two paths in a graph are called *compatible* if they both start in the same vertex and both end in the same vertex. Given compatible paths *P* and Q\** and a path *P* that contains *P\** as a subpath, the (*P\**,*Q\**)-*rerouting* substitutes *P* by a new path where the subpath *P\** of *P* is substituted by *Q*.* Given a low-coverage subpath *P\** of a read-path *P*, mowerDBG attempts to find a read-path containing a higher-coverage *P\**-compatible subpath *Q*.* Afterward, it corrects errors in *P* by performing the (*P\**,*Q\**)-rerouting of *P* and reduces (increases) the coverage of all edges in the subpath *P\** (*Q\**) by 1.

We denote the edit distance between strings *v* and *w* as *distance*(*v,w*). mowerDBG classifies strings *v* and *w* as *similar* if *distance*(*v,w*) ≤ *fraction*\*min{|*v*|,|*w*|} and *possibly-similar* if the difference between their lengths does not exceed *fraction*\*min{|*v*|,|*w*|} (*fraction* is a parameter with the default value=0.01). Two compatible paths in the de Bruijn graph are *similar* (*possibly*-*similar*) if they spell similar (possibly-similar) strings.

Given compatible and similar subpaths *P\** and *Q\** of two read-paths in the de Bruijn graph, *Q\** is classified as a *bypass of P\** if it is a high-coverage subpath. For each low-coverage internal subpath *P\** of a read-path *P,* mowerDBG searches for a read-path *Q* and its subpath *Q\** that represents a bypass of *P\**. It classifies a low-coverage internal subpath as a *no-bypass, uni-bypass,* or *multi-bypass* if it has no bypasses, a single bypass, or multiple bypasses, respectively.

Since classifying bypasses into these categories may be time-consuming (particularly, in the case of long subpaths), mowerDBG uses a slightly different but fast classification of external bypasses. Specifically, for each low-coverage internal subpath *P\** connecting vertices *source* and *sink*, mowerDBG considers all read-paths that traverse both *source* and *sink* and identifies all possibly-similar subpaths of these read-paths that are compatible with *P\**. If there are no high-coverage subpaths in this set, *P\** is classified as a no-bypass. If all high-coverage subpaths in this set are identical (resulting in a subpath *Q\**), *P\** is classified as a uni-bypass. Otherwise, it is classified as a multi-bypass.

There are 416157 (79398) low-coverage internal (external) subpaths in the graph *DB*(T2TX,501). 444, 414,399, and 1,314 out of 416157 low-coverage internal subpaths are no-bypasses, uni-bypasses, and multi-bypasses, respectively. Below we describe how mowerDBG reroutes uni-bypasses and multi-bypasses (no-bypasses are not affected by error correction).

#### Rerouting a uni-bypass

Since a uni-bypass *P\** has a single bypass *Q\**, it is a candidate for a (*P\**,*Q\**)-rerouting if *P\** and *Q\** are similar. However, mowerDBG skips the time-consuming similarity check since possibly-similar subpaths turned out to be similar in the vast majority of cases. After rerouting all internal uni-bypasses, 76% of reads in the T2TX dataset become error-free. Only 1522 out of 414,399 internal uni-bypasses (0.35%) in the graph *DB*(T2TX,501) resulted in invalid re-routings. We note that an invalid re-routing does not necessarily lead to an error in the final assembly since it is typically corrected at the follow-up error-correction steps. mowerDBG reroutes all uni-bypasses in reads before starting a more complex multi-bypass rerouting step.

#### Rerouting a multi-bypass by an iterative analysis of read-paths

Error-correcting a read sampled from a repetitive region is challenging since it often requires deciding between multiple rerouting options. For example, a repeat with many slightly different copies forms a “cycle with many bulges” that is traversed slightly differently by different repeat copies, resulting in multiple correct paths between the same vertices. mowerDBG uses two techniques to reroute a multi-bypass: (i) iterative analysis of read-paths through the graph and (ii) identifying a bypass that is most similar to a low-coverage subpath.

To use information about read-paths through the graph, mowerDBG considers longer *k^+^*-mers (for *k^+^ > k*) and assumes that nearly all genomic *k^+^*-mers are covered by at least *minKmerCoverage* reads (default value is 4). Let *Q\** be a bypass of *P\** such that *Q\** is a subpath of *Q* and *P\** is a subpath of *P.* A subpath *Q_1_Q*Q_2_* of *Q* is called an *extension* of the bypass *Q\** by a read-path *Q* if *Q_1_P*Q_2_* is also a subpath of *P*, i.e, *Q_1_Q*Q_2_* coincides with a subpath of *P* everywhere except for *P\**. We denote the nucleotide length of the longest extension of the bypass *Q\** by a read-path *Q* as *extension*(*Q*,Q*).

Given a value *k^+^,* we say that a read *Q supports* a bypass *Q\** if *extension*(*Q*,Q*) ≥ *k^+^* and classify a bypass as *well-supported* if it is supported by at least *minKmerCoverage* reads. mowerDBG applies the (*P*,Q\**)-rerouting to the path *P* if (i) *P*\* has only one well-supported bypass *Q\**, and (ii) all other bypasses are supported by a single read. mowerDBG uses the default sequence *k^+^*=800, 2000, 3500 to perform iterative rerouting with the gradually increasing values of *k^+^*. In the case when there are multiple well-supported bypasses for the final value of *k^+^* (a common situation in highly-repetitive regions, resulting in 2035 remaining multi-bypasses after this step), mowerDBG selects one of them based on analyzing the edit distances as described below.

#### Rerouting a multi-bypass by identifying the most similar bypass

Given a multi-bypass *P\**, mowerDBG computes its edit distance with each its bypass to identify the most similar bypass *Q*_sim_*. Although (*P\**,*Q*_sim_*)-rerouting corrects errors in *P\** in the vast majority of cases, in rare cases it introduces an error. To prevent such errors, mowerDBG performs a *triangle test* inspired by a similar test in the mosaicFlye assembler (Bankevich and Pevzner, 2020). For each bypass *Q\** of *P\** (that is different from *Q*_sim_*), it tests if *distance*(*P*,Q\**)*=distance*(*P*,Q*_sim_*)*+distance*(*Q*_sim_,Q*).* If this test holds, mowerDBG performs the (*P\**,*Q*_sim_*)-rerouting of the multi-bypass *P\**. In most cases, the triangle test is equivalent to checking whether the edit operations to transform *P\** into *Q*_sim_* represent a subset of edit operations to transform *P\** into *Q*,* a rather strong condition. The triangle test leads to correcting 1790 out of 2035 remaining multi-bypasses (1511 of them represent valid corrections).

#### Rerouting an external bypass

Supplementary Figure S10 illustrates that simply deleting all low-coverage external subpaths may significantly reduce the length of some reads and thus negatively affects the ability of multiplexDBG to resolve repeats. Given a low-coverage external subpath *P\** of a read-path *P*, mowerDBG thus attempts to error-correct *P* by substituting *P\** with a high-coverage subpath in *DB*(*Reads,k*).

Below we limit attention to *suffix-subpaths* ending in the last edge of a read (external *prefix-subpaths* are analyzed similarly). Given a suffix-subpath *P\**, mowerDBG considers all reads passing through its first vertex and identifies their high-coverage subpaths that start at this vertex. For each such subpath, it identifies its prefix that has the lowest edit distance from *P\** and analyzes the set of such prefixes. Similar to the classification of each internal low-coverage subpath into a no-bypass, uni-bypass, or multi-bypass, mowerDBG classifies each low-coverage suffix-subpath into a *no-suffix*, *uni-suffix*, or *multi-suffix.* It applies the previously described rerouting algorithm for three types of bypasses but this time applied to the three types of suffix-subpaths.

This procedure reroutes 79150 out of 79398 (99.7%) low-coverage external subpaths. mowerDBG removes the remaining 248 low-coverage external subpaths, thus shortening the corresponding reads.

#### Bulge-collapsing

We refer to all parallel edges between vertices *v* and *w* in the graph *DB*(*Reads,k*) as a *bulge*. An edge in a bulge with the highest coverage is classified as a *heavy* edge (a bulge may have multiple heavy edges) and all other edges are classified as *light edges*.

We classify a bulge as *collapsible* if the total coverage of its edges does not exceed *coverageAmplifier*・*medianCoverage*, where *coverageAmplifier* is a parameter (the default value 1.5) and *medianCoverage* stands for the median coverage of all (*k*+1)-mers in the graph. An edge in a bulge is *correct* if it represents a substring of the genome and *erroneous*, otherwise. A collapsible bulge is *reducible* if it has a single correct edge and *foolproof* if this single edge is heavy.

Bulge-collapsing removes all light edges in a bulge and reroutes all read-paths containing these edges through a heavy edge of this bulge (if a bulge has multiple heavy edges, an arbitrary heavy edge is chosen). The previously described path-rerouting step results in the de Bruijn graph *DB*(T2TX_reroute_,501) on the error-corrected read-set T2TX_reroute_ that has 144 bulges (140 of them are collapsible). Moreover, 128 (124) out of 140 collapsible bulges in *DB*(*T2TX*_reroute_,501) are reducible (foolproof). Since the vast majority of collapsible bulges in this graph are foolproof, mowerDBG collapses all collapsible bulges until no such bulges are left.

#### Correcting dimers

The second most common type of errors in HiFi reads (after errors in the length of homopolymer runs) is manifested as incorrect length of long *dipolymer runs*, i.e., regions formed by only two nucleotides. Since LJA collapses homopolymers in reads (and later expands the collapsed homopolymers in HPC contigs with LJApolish), it collapses each dipolymer run into a tandem *dimer repeat*.

Different HPC reads covering a long dimer repeat often represent this repeat with differing copy numbers. For example, some reads may represent a dimer repeat …AGAGAG… with the correct copy number 3 while others may represent it as …AGAGAGAG… with incorrect copy number 4. Such errors are common in *long dimer repeats* with copy number at least *longDimerRepeat* (default value 16). Moreover, the number of reads with incorrect copy number may exceed the number of reads with correct copy number in some cases. Instead of trying to guess the correct copy number, mowerDBG corrects all reads covering a dimer repeat in such a way that they have the same (possibly incorrect) copy number. It thus considers all dimer repeats in a read (including short dimer repeats of multiplicity 2), finds out the most “popular” copy number among all reads covering this repeat, and uses it to correct (or corrupt) all such reads in a consistent way. In addition, it collapses all long dimer repeats in reads into repeats of multiplicity exactly *longDimerRepeat*. The correct sequences of the corrupted dimer repeats are later reconstructed by the LJApolish module.

#### Correcting pseudo-correct reads

The error-correction procedures described above assume reads formed by high-coverage (*k+*1)-mers are correct. However, there exist *pseudo-correct* reads that do not represent a substring of the genome even though they are formed by high-coverage (*k*+1)-mers. For example, an error in a read sampled from a repeat may result in a (*k*+1)-mer that coincides with a (*k*+1)-mer from a different copy of the same repeat, resulting in a pseudo-correct read. Identifying pseudo-correct reads and correcting them is a challenging task that LJA addresses only at the second round of error correction with a larger *K*-mer size.

We denote the set of reads after the first three error-correcting steps (path-rerouting, bulge-collapsing, and dimer-correcting) in the graph DB(T2T’,5001) as T2T’’. Below we describe how mowerDBG identifies and error-corrects 71647 pseudo-correct reads in this read-set.

To identify pseudo-correct reads, mowerDBG approximates the set of *unrepeated* edges, i.e., edges in the de Bruijn graph that are visited exactly once by the genome traversal. It considers the set of all *ultralong* edges of length exceeding the *maxRepeatLength* threshold (default value 40 kb) as the first approximation for the set of unrepeated edges. This large threshold is chosen to ensure that all (or the vast majority of) perfect repeats in the genome have length below *maxRepeatLength*, implying that all (or nearly all) ultralong edges are unrepeated (Supplementary Note 3). We refer to the resulting edges as *unique* and acknowledge that this procedure may fail to classify some unrepeated edges as unique. 5090 out 23010 edges in the de Bruijn graph *DB*(T2T’’,5001) are unique. Supplementary Note 9 describes how mowerDBG extends the set of the originally selected unique edges (from 5090 to 13562) by finding maximum flows in the specially constructed networks.

For each unique edge *e,* mowerDBG identifies the set *Paths*(*e*) of all read-paths containing this edge. We denote the set of their *suffix-paths* starting at the end of *e* as *Suffixes*(*e*) and the set of their *prefix-paths* ending at the start of *e* as *Prefixes*(*e*). We classify two paths in a graph as *consistent* if one of them is a subpath of another, and *inconsistent* otherwise. A suffix-path (prefix-path) that contains most other paths from *Suffixes*(*e*) (*Prefixes*(*e*)) as subpaths is called a *representative* suffix-path (prefix-path) for the edge *e*. We define a *representative* path for the set *Paths*(*e*) as the path formed by the representative prefix-path, followed by the edge *e*, and followed by the representative suffix-path.

Since all error-free read-paths through a unique edge *e* are consistent, a path-set *Path*(*e*), that contains inconsistent paths, must contain some paths originating from pseudo-correct reads. To correct such reads, mowerDBG considers each read-path in *Path*(*e*) that is inconsistent with the representative path of *Paths*(*e*) and substitutes it by the representative path. While this procedure, that corrects 71647 pseudo-correct reads in the T2T’’ dataset. has only a small impact on the overall error rate in reads, it is important since each pseudo-correct read may trigger a misassembly. In addition, this procedure corrects bulges caused by pseudo-heterozygosity in the CHM13 cell line.

#### Two-round error-correction

After read-rerouting, bulge-collapsing, and dimer-correcting in the graph *DB*(*T2T,*501), 92.5% of reads in the error-corrected read-set have perfect alignment to the HPC reference genome and the error rate in error corrected read-set T2T’ is reduced to 3.8 errors per megabase of the total read-length. The remaining erroneous *k*-mers are often supported by multiple reads and thus are difficult to distinguish from variations between repeat copies (for *k*=501).

To further reduce the error rate, LJA launches jumboDBG to construct *DB*(*T2T’,K*) and performs the second error-correcting round with larger *K*-mer size (default value *K*=5001) using this graph. This two-round approach results in an even more accurate read-set T2T*. We note that the last step of mowerDBG (correcting pseudo-correct reads) is invoked only at the second round of error correction.

The rationale for the second error-correcting round is that it becomes less likely for the same error to be supported by *K*-mers than by *k*-mers. Indeed, the graph *DB*(*T2TX’,*5001) has 750 bulges and 744 of them are collapsible. 738 (536) out of 744 collapsible bulges in *DB*(*T2TX’,*5001) are reducible (foolproof) bulges. After correcting 7990 reads using the path-rerouting procedure with large *K*-mer size (5001), we are left with 484 no-bypasses and no multi-bypasses. The remaining external and internal no-bypasses are then removed from the graph by cutting off the ends of the reads with external bypasses and splitting reads with internal bypasses into several shorter reads.

After two error-correcting rounds, the initial T2TX read-set is transformed into the error-corrected read-set T2TX* with error rate in reads reduced to only 2.4 errors per megabase and only 2.6% of reads have errors. Moreover, the de Bruijn graph constructed from error-corrected reads for *k*=5001 does not contain low-coverage edges, suggesting that the remaining 2.6% of reads are corrected consistently with each other even though they do not perfectly match the reference genome due to either heterozygous sites or the repeat-induced read corruption that refers to corrupting reads that span one repeat copy to match another, slightly different copy. We emphasize that the repeat-induced read corruption does not necessarily lead to errors in the final assembly since the LJApolish module fixes nearly all errors in corrupted reads.

After two error-correcting rounds, mowerDBG identifies all reads that still contain uncorrected bypasses (1395 reads for chrX) and breaks each such read into shorter reads by removing each uncorrected bypass. After all error-correction steps, the compressed de Bruijn graph of the error-corrected T2TX dataset on 5001-mers consists of 154 vertices and 223 edges and is broken into 3 connected components, indicating breaks in coverage of 5001-mers.

### Supplementary Note 9: Using flows in networks to identify unrepeated edges

When the genome is known, we define the *flow* through an edge *e* in its compressed de Bruijn graph (denoted as *flow*(*e*)) as the number of times this edge is visited by the genome traversal. Given a vertex *v*, we define *inflow*(*v*) (*outflow*(*v*)) as the sum of flows through all incoming edges in *v* (outgoing edges from *v*). The flow is *balanced* (i.e., *inflow*(*v*)=*outflow*(*v*)) for all vertices in *DB*(*Genome,k*) except for possibly *k*-prefixes and *k*-suffixes of linear chromosomes.

When the genome is unknown, our goal is to estimate the flow through all edges in *DB*(*Reads,k*). Although this problem was addressed by Pevzner and Tang, 2001 (for finding lower bounds for flows) and Nurk et al., 2013 (for finding chimeric edges with zero flow), we are addressing a slightly different problem of finding unrepeated edges with flow 1.

Given a graph with an identified subset of unique edges *U* (initially, all ultralong edges), mowerDBG iteratively extends it as follows. It first “breaks” each edge (*v,w*) in *U* by removing this edge and adding two new vertices *v\** and *w\** along with edges (*v,v\**) and (*w*,w*). Afterward, it considers each connected component in the resulting graph, “glues” all newly added vertices in this component into a single *nick* vertex, and classifies all edges incident to the nick vertex as *nick* edges. Since the genome traversal induces an (unknown) traversal of each resulting connected component and since each edge from *U* is unique, the nick vertex is balanced and each nick edge is traversed once (has flow equal to 1). The traversal of each component induces an (unknown) balanced flow except for (rare) cases when a component contains a *k*-prefix or a *k*-suffixes of a linear chromosome.

Since the flows are unknown (except for nick edges), it is not immediately clear how to find additional unique edges in each connected component. However, if the flow through a non-nick edge (*s,t*) in this component is equal to 1 for *all* possible balanced flows, this edge is unique. To estimate the maximum flow through this edge, mowerDBG assigns capacity 1 to all nick edges, capacity ∞ to all other edges, and finds the maximum flow (satisfying the capacity constraints) from *t* to *s* in the resulting network. An edge (*s,t*) is classified as unique if the maximum flow from *t* to *s* is 1. mowerDBG iteratively adds all newly identified unique edges to the set of the previously identified unique edges until no new unique edges are identified. This algorithm identifies 13562 unique edges (out of a total of 23010 edges) in the graph *DB*(*T2T’’*,5001) on the error-corrected read-set *T2T’’*.

### Supplementary Note 10: Fast graph transformation algorithm

Let *Paths* be the set of all read-paths in the compressed de Bruijn graph *G=DB*(*Reads*,*k*). A straightforward approach to constructing the graph *G*(*Paths*) nearly doubles the path lengths at each iteration and thus faces the time/memory bottleneck. However, the compressed de Bruijn graph is getting less tangled with an increase in the *k-*mer size, implying that the vast majority of the newly introduced red edges are merely subpartitions of longer non-branching paths. Below we describe how multiplexDBG avoids this time/memory bottleneck.

The path-graph *G*(*Paths*) is a subpartition of the graph *DB*(*Reads*,*k*+1) (after properly defining edge-labels and ignoring colors of edges). For each edge *e* in the graph *DB*(*Reads,k*), we maintain the set of paths *Paths*(*e*) containing this edge. A path *e_1_, e_2_, e_3_*,… in *DB*(*Reads*, *k*) corresponds to a blue-red path *e_1_,* transition edge between *e_1_* and *e_2_*, *e_2_,* transition edge between *e_2_* and *e_3_*, *e_3_*,… in *G*(*Paths*), where labels of blue edges have lengths at least (*k*+1) and labels of red edges represent (*k*+2)-mers. Therefore, a straightforward approach to constructing the graph *G*(*Paths*), that recomputes labels from scratch at each iteration, nearly doubles the path lengths at each iteration and thus faces the time/memory bottleneck.

Given a path-set *Paths* in a graph *G*, we call edges (*v,w*) and (*w,u*) in *G paired* if the transition-set *Transitions*(*Paths)* contain this pair of edges. A vertex *w* in *G* is *paired* if each edge incident to *w* is paired with at least one other edge incident to *w*, and *unpaired*, otherwise. For a simple paired vertex, the local topology of the graph “around” this vertex remains the same after the graph transformation. In the framework of the read-paths in the multiplex de Bruijn graph (when the read-set is complemented by virtual reads), the local topology of both paired and unpaired simple vertices (with the exception of the dead-end simple vertices) remains the same after this transformation. The transformation of a complex *N*-in-*M*-out vertex *w* results in substituting this vertex by two sets of vertices (one of size *N* and another of size *M*) and adding up to *N* ・ *M* red edges (that connect vertices from these two sets) to the graph. Each path from *Paths*(*w*) traversing *t* complex vertices will be transformed into a path with *t* red edges and the non-branching paths formed after this transformation have to be transformed into single edges. Below we describe how multiplexDBG speeds-up graph transformations.

For each parameter *k*, the vast majority of vertices in the compressed de Bruijn graph of reads are 2-in-1-out and 1-in-2-out vertices. Below we consider the graph transformation of a 2-in-1-out vertex *w* with incoming edges *in_1_* and *in_2_* and the outgoing edge *out* (transformations of *N*-in-1-out and 1-in-*N*-out vertices are performed similarly). This transformation merely substitutes the *k*-mer label of this vertex by the (*k*+1)-mer *label*(*w*)*\*symbol_k+1_*(*out*). Since it preserves the label of the edge *out* and adds a single symbol *symbol_k+1_*(*out*) after the end of labels of edges *in_1_* and *in_2_* (Figure 5.A), multiplexDBG performs fast graph transformation by simply adding a single symbol to the labels of incoming edges.

To transform a complex *N*-in-*M*-out vertex *w* (Figure 5.B), multiplexDBG generates the list of paths traversing each new red edge (for up to new *N*M* red edges for each complex *N*-in-*M*-out vertex) using the list of paths *Paths*(*v,w*) traversing each edge (*v,w*) in the graph. We define *Paths^+^*(*v*) as the set of all paths visiting incoming edges into vertex *v* (each path in *Paths^+^*(*v*) either traverses *v* or stops at *v*) and note that the transformation of a complex vertex *w* takes |*Paths^+^*(*w*)| time. The number of operations to transform all complex vertices is bounded by the sum of |*Paths^+^*(*v*)| over all complex vertices in the graph. Since the number of complex vertices in the compressed de Bruijn graph is small and since processing a simple vertex takes constant time, the transformation of *DB*(*Reads*, *k*) into *DB*(*Reads*, *k*+1) is fast for large *k* even though it can be rather slow for small *k*. For example, constructing the graph *DB*(chrX,501) using jumboDBG and further transforming it into the graph *DB*(chrX,5001) using graph transformations takes 5+393 minutes, while the direct construction of *DB*(chrX,5001) using jumboDBG takes just 3 minutes. However, iterative increasing of the *k*-mer size during the construction of the multiplex de Bruijn graph is only crucial for large *k* (e.g., greater than 5001) that can be done rapidly.

### Supplementary Note 11: Limitations of the multiplex de Bruijn graph transformation

Boucher et al., 2015 described the *variable-order de Bruijn graph* that compactly represents information about the de Bruijn graph of a read-set across multiple *k*-mer sizes. Lin and Pevzner, 2014 described a theoretical approach for constructing the de Bruijn graphs with multiple *k*-mer sizes that however was not designed for practical genome assembly challenges. The multiplex de Brujin graph algorithm differs from these approaches since it incorporates vertices with varying *k*-mer-sizes and greatly improves the contiguity of HiFi assemblies.

We refer to a read-set as *incomplete* if it does not contain reads supporting some genomic transitions through a vertex in the de Bruijn graph. Similarly to the iterative de Bruijn graph approach (Peng et al., 2010), although the graph *MDB*(*Reads,k*) results in a more contiguous assembly than *DB*(*Reads,k*), there is a risk that some multiplex graph transformation may “destroy” the genome traversal and even lead to assembly errors in the case of an incomplete read-set. Below we describe these risks and illustrate that multiplex transformations may be overly-optimistic (by transforming vertices that should have been frozen) and overly-pessimistic (by freezing vertices that should have been transformed).

Supplementary Figure S11 shows a circular genome ARDARCBRCE that traverses the repeat R three times via subpaths ARD, ARC, and BRC and an incomplete read-set that supports only two of these subpaths (ARD and BRC). This example illustrates the case when a graph transformation results in a fragmented assembly since the multiplex de Bruijn graph of reads “loses’ ’ the genome traversal. Indeed, after a series of transformations, when the *k*-mer size becomes equal to the length of the repeat R, this repeat will be transformed into a single “red” vertex that will be classified as paired because each incoming edge into this vertex is paired with an outgoing edge from this vertex (and vice versa).

**Supplementary Figure S11.**
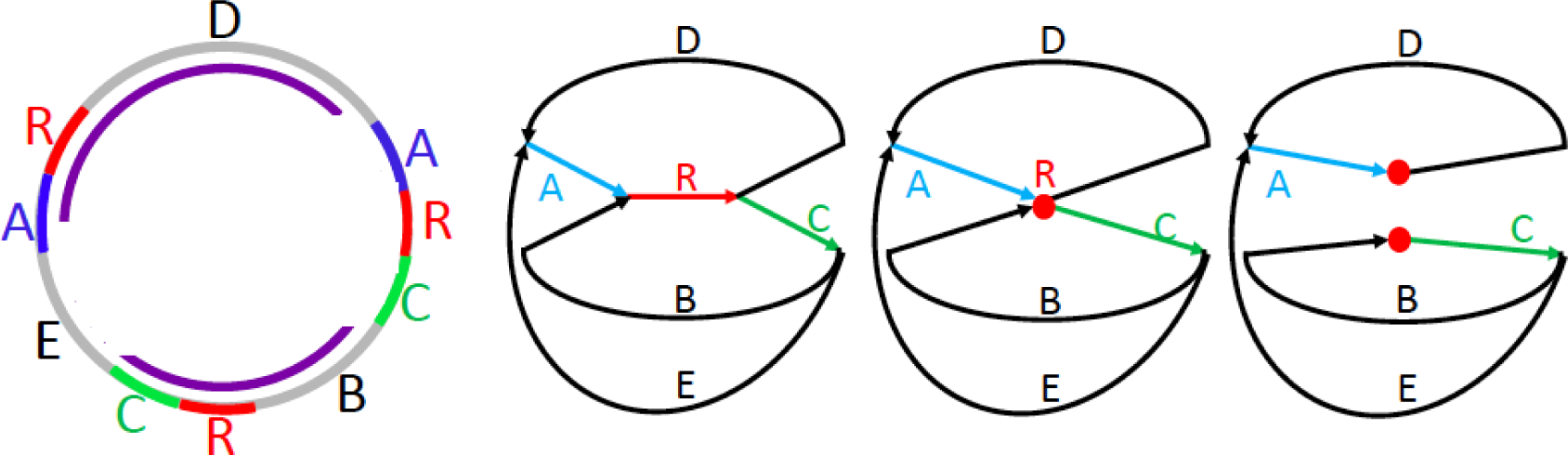
A circular genome ARDARCBRCE (first panel), its compressed de Bruijn graph (second panel), its multiplexed de Bruijn graph after the edge R is transformed into a paired vertex (third panel), and its multiplexed de Bruijn graph after transforming this paired vertex (fourth panel). The read-set includes each two-edge path corresponding to a pair of consecutive edges in the genome as well as the three-edge paths ARD and BRC shown in purple. However, it does not include the three-edge path ARC. After the repeat R is transformed into a single vertex, this vertex is classified as paired since each incoming edge is paired with an outgoing edge and vice versa (A is paired with D and B is paired with C). Transforming this vertex results in a multiplex de Bruijn graph that does not adequately represent the genome since it “loses” the genome traversal.

Another limitation of the multiplex graph transformation is illustrated by an example of a linear genome ARBRC and an incomplete read-set that contains a read ARB spanning one instance of the repeat R but does not contain a read BRC spanning another instance of this repeat. In this case, after a series of transformations, when the *k*-mer size becomes equal to the length of the repeat R, this repeat will be transformed into an unpaired vertex that has to be frozen. However, an addition of an *auxiliary read* BRC (when it is relatively “safe”) reclassifies this vertex as paired and enables a graph transformation at this vertex.

### Supplementary Note 12: LJApolish algorithm for expanding collapsed assemblie

#### Expanding error-free HPC contigs using error-free reads

We define the *run-length* of a nucleotide (position) in an HPC read as the length of the homopolymer run in the original read that was collapsed into this nucleotide. If original (uncollapsed) reads and HPC contigs were error-free, one would be able to transform each original read into an error-free HPC read and align it against an HPC contig using exact pattern matching. We refer to each pair of aligned positions in an HPC reads and an HPC contig as *aligned positions*. Given all pairs of aligned positions, the contig expansion problem has a simple solution: one can simply consider all positions in HPC reads aligned to a given position *pos* in an HPC contig and define its run-length *run*(*pos*) as the (same) run-length of all positions in reads aligned to *pos*. Substituting a nucleotide at position *pos* in an HPC contig by a run of this nucleotide of length *run*(*pos*) would generate an error-free (uncollapsed) contig.

Even when original (uncollapsed) reads have errors but all HPC reads and HPC contigs are error-free, one can apply a similar approach to accurately expand HPC contigs. Indeed, after transforming each original error-prone read into an error-free HPC read and aligning it against an error-free HPC contig (again using exact pattern matching), one can examine all positions in reads that are aligned against a position *pos* in an HPC contig. However, since original reads are error-prone, aligned positions in different reads may have different run-lengths. We therefore define *median-run*(*pos*) as the median run-length of all positions in reads aligned to *pos*. Since error-rate in HiFi reads is low, substituting a nucleotide at position *pos* in an HPC contig by a run of this nucleotide of length *median-run*(*pos*) would generate an (uncollapsed) contig with low error rate.

In reality, since both HPC reads and HPC contigs are error-prone, one needs an extremely accurate and fast algorithm for their alignment and follow-up expansion of HPC contigs, a challenging task for reads from highly-repetitive regions.

#### Mapping reads to the compressed de Bruijn graph

Since mapping reads to highly-repetitive regions is a challenging task (Jain et al., 2020, Mikheenko et al., 2020), LJApolish uses information about the read-path of each read that jumboDBG generates after constructing the de Bruijn graph.

After constructing *DB*(*Reads,k*), jumboDBG finds the read-path of each read through this graph and identifies the starting (ending) position of this read within the starting (ending) edge of its read-path. To address this problem for a given edge, jumboDBG stores all *k*-mers starting at positions 0, *width,* 2 ・ *width*, 3 ・ *width*, etc. in the label of this edge (*padded k-mers)*. Since it discards all reads shorter than *width+k*-1 to ensure that each remaining read contains at least one minimizer, every read contains at least one padded *k*-mer. jumboDBG combines all padded *k*-mers for all edges and all junctions in *DB*(*Reads,k*) into a single set and traverses all *k*-mers in each read to find out how this read traverses all vertices of this combined set. Using this information, it generates the read-path for each read and identifies the starting/ ending position of each read within the starting/ending edge of a read-path.

#### Aligning error-corrected HPC reads to HPC contigs.]

LJA reports labels of edges of the constructed de Bruijn graph as HPC contigs. LJApolish expands them by aligning each HPC reads against the corresponding HPC contig. To facilitate this task, LJA provides LJApolish with a “hint” for the starting/ending locations of the alignment of each HPC read against the corresponding HPC contig.

Since LJA error-corrects both HPC reads and the de Bruijn graph in a coordinated fashion, each error-corrected HPC read (or its segment) represents a substring of an HPC contig. Thus, the alignment of an *error-corrected* HPC read to HPC contigs amounts to a simple problem of finding an exact match. As described above, LJA finds this exact match and uses its starting/ending locations as a hint for finding accurate alignments of the original (uncollapsed) read against HPC contigs.

However, this description does not take into account that some HPC reads were trimmed and even broken into subreads by mowerDBG. Thus, the locations of alignments of the error-corrected HPC reads represent approximate locations of alignments of the original HPC reads to the HPC contigs, further complicating construction of the nucleotide-by-nucleotide alignment based on the information about locations. LJApolish uses the provided locations to rapidly construct a nucleotide-by-nucleotide alignment between HPC reads and HPC contigs.

#### Banded alignment of HPC reads to HPC contigs

LJA provided the starting/ending locations of alignments of error-corrected HPC reads against HPC contigs for all 5,567,158 reads in the T2T dataset, except for 5912 reads (0.1%) it was unable to correct. As an input, LJApolish takes the assembled HPC contigs, the original (uncollapsed) HiFi reads, and the locations of the alignments of the error-corrected HPC reads to the HPC contigs provided by LJA. Alternatively, one can use read mapping tools, such as minimap2 (Li, 2018), to generate locations without the hints provided by LJA and thus potentially use LJApolish for polishing contigs generated by other assemblers. However, since read mapping in highly-repetitive regions is a difficult problem (Mikheenko et al., 2020, Jain et al., 2020), this approach may result in an inferior accuracy with respect to single-base errors.

To align an HPC read against the corresponding fragment of an HPC contig within the specified starting/ending locations. LJApolish constructs a (global) *banded alignment* in *O*(*LE*) time, where *L* is a read of length *L* and *E* is the number of errors in this read. Since HiFi reads are accurate, this approach represents a speedup as compared to the prohibitively slow standard alignment approach with *O*(*L^2^*) running time. Since collapsing all homopolymer runs and all long dinucleotide runs in a read typically results in an HPC read with a small number of errors (Supplementary Note 8), LJApolish computes the banded alignments of each HPC read against the corresponding fragment of an HPC contig. To construct such banded alignment, it uses the ksw2 library (https://github.com/lh3/ksw2) with the default *bandwidth* equal to 10 but if the alignment goes out of band, the *bandwidth* threshold is iteratively doubled until the alignment is constructed within a specified band. To limit the time for aligning a single read, this iterative doubling process stops when the bandwidth reached *bandwidthLarge* (default value *bandwidthLarge*=320). Only 1421 out of 5,561,246 reads (0.03%) with specified locations failed to align within the maximum bandwidth.

#### Expanding HPC contigs

For each pair of aligned positions in the constructed read-contig alignment, LJApolish stores the run-length of this position in the HPC read. For each position in the HPC contig, it stores at most first *max_coverage* run-lengths of the aligned positions in HPC reads to limit the memory consumption (default value *max_coverage=*20).

Given the set of run-lengths of HPC reads covering a given position in an HPC contig, LJApolish computes the median of run-lengths of the corresponding positions in reads and expands the nucleotide at this position into a homopolymer run of length equal to this median. The median (rather than the average) is selected to reduce the influence of outliers with incorrect run-lengths in reads.

#### Analyzing dinucleotide runs

Since dinucleotide runs in HiFi reads are often corrupted (e.g., …CATATATG… may be represented as …CATATATATG…), they may trigger assembly errors, often requiring an additional *dinucleotide correction step* (Nurk et al., 2020, Cheng et al., 2021). Such errors in the original HiFi reads may propagate into HPC reads and “shift” the alignments of reads, particularly in regions with long dinucleotide runs. There exists 307702 *long* dinucleotide runs in the T2T assembly with multiplicities at least *long-direpeat* (default value 10).

To minimize this effect, LJApolish marks all long dinucleotide runs in each HPC contig prior to the read alignment procedure. For each read-alignment covering a marked region, it extracts the fragment of the (uncollapsed) HiFi read that corresponds to this region and stores such fragments separately. Afterwards, instead of selecting the median multiplicity for each position in the compressed contig, LJApolish calculates the consensus string for these fragments by constructing their *partial order alignment* (Lee et al, 2002). It uses the *spoa* library that outputs the heaviest path in the partial order graph as the consensus string (https://github.com/rvaser/spoa).

#### Analyzing single-base errors

We analyzed all 34259 mismatches and indels in the homopolymer-collapsed LJA assembly of the T2T read-set, reported by QUAST-LG that was updated to incorporate the latest 2.21 version of minimap2 version and to include a report of all mismatches and indels. 4939 errors (11%) occur in the low-coverage regions (coverage below 5x). 2009 of these errors (6%) occur within long dinucleotide runs (with multiplicity at least 10) and 31 (0.06%) occur within long homopolymer runs (with length at least 10 bp). Additional 6201 errors (18%) occur in long dinucleotide runs with coverage at least 5x. Therefore, low-coverage regions and long dinucleotide runs account for 29% of all errors.

Outside the low-coverage regions and dinucleotide runs, long homopolymer runs (with length at least 10 bp) account for 13505 (39%) errors. (Supplementary Table S3). Complex indels, formed by more than two different nucleotides, that are inherited from the compressed LJA assembly account for 746 (2%) errors. The remaining 9868 (29%) errors are difficult to classify as they may be caused by a variety of reasons: heterozygosity of the CHM13 cell line, collapsed inexact repeats in the assembly, non-optimal selection of alignment parameters in QUAST-LG, etc.

**Supplementary Table S3.** Distribution of the number of errors over the homopolymer runs of various lengths. All homopolymer runs in the T2T assembly are shorter than 90 bp.

| Length of the homopolymer run | Number (percentage) of errors in the LJA assembly | Total number of homopolymer runs |
| --- | --- | --- |
| 10-15 | 1157 (0.19%) | 610840 |
| 15-20 | 1685 (0.73%) | 232006 |
| 20-25 | 3024 (2.94%) | 102990 |
| 25-30 | 4273 (10.66%) | 40074 |
| 30-35 | 1772 (18.64%) | 9508 |
| 35-40 | 843 (22.19%) | 3799 |
| 40-45 | 465 (28.66%) | 1622 |
| 45-50 | 168 (25.07%) | 670 |
| 50-55 | 87 (29.19%) | 298 |
| 55-60 | 25 (26.88%) | 93 |
| 60 and longer | 6 (24.00%) | 25 |

#### QUAST evaluation of the uncollapsed T2T assembly

Since QUAST inflates the number of misassemblies, we evaluated them by running QUAST on homopolymer-collapsed contigs and using the homopolymer-collapsed *T2TGenome* as a reference. Supplementary Table S4 provides QUAST results without homopolymer-collapsing.

**Supplementary Table S4.**
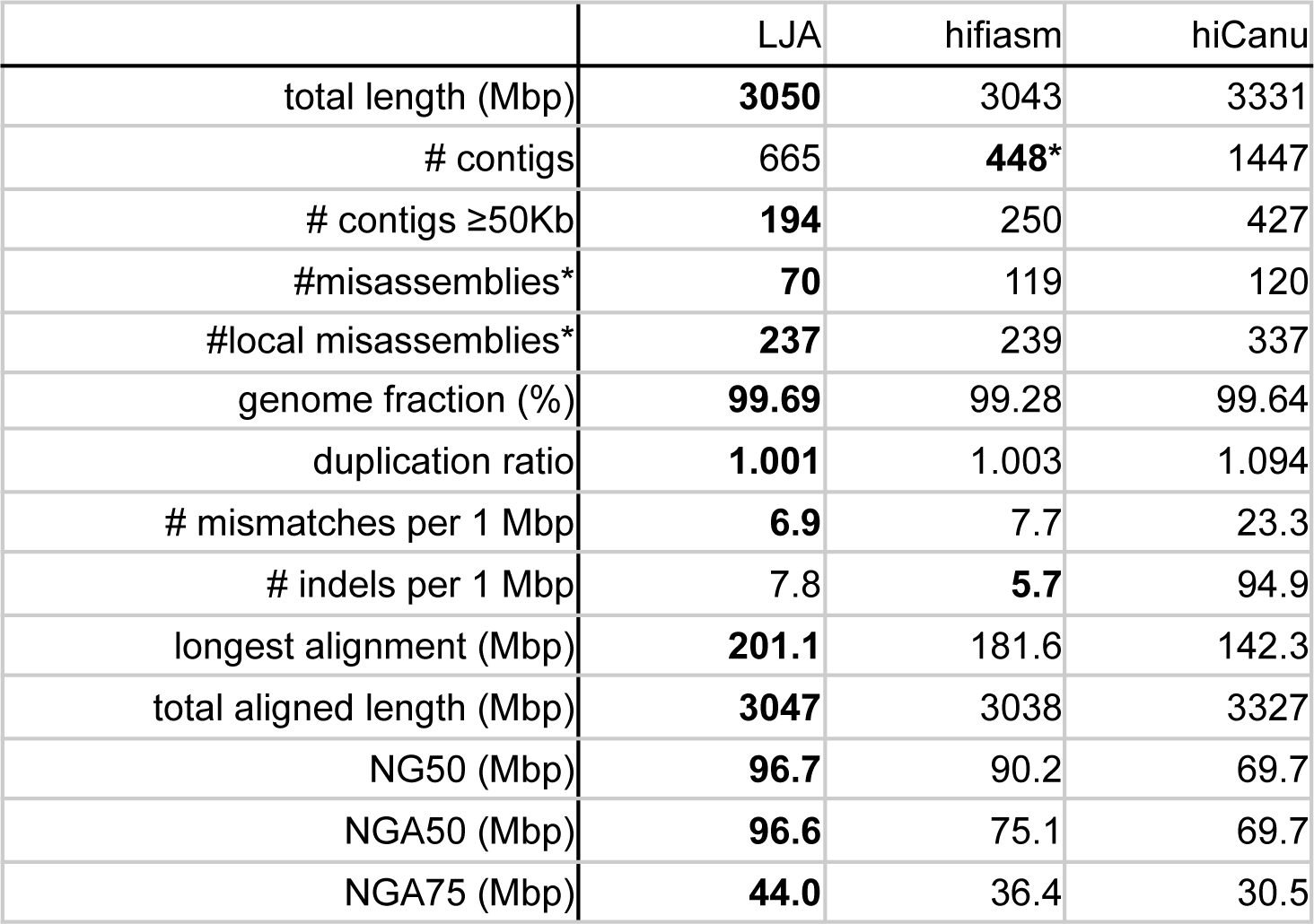
Benchmarking LJA, hifiasm, and HiCanu on the T2T dataset without homopolymer-collapsing.

|  | LJA | hifiasm | hiCanu |
| --- | --- | --- | --- |
| total length (Mbp) | **3050** | 3043 | 3331 |
| # contigs | 665 | **448\*** | 1447 |
| # contigs ≥50Kb | **194** | 250 | 427 |
| #misassemblies* | **70** | 119 | 120 |
| #local misassemblies* | **237** | 239 | 337 |
| genome fraction (%) | **99.69** | 99.28 | 99.64 |
| duplication ratio | **1.001** | 1.003 | 1.094 |
| # mismatches per 1 Mbp | **6.9** | 7.7 | 23.3 |
| # indels per 1 Mbp | 7.8 | **5.7** | 94.9 |
| longest alignment (Mbp) | **201.1** | 181.6 | 142.3 |
| total aligned length (Mbp) | **3047** | 3038 | 3327 |
| NG50 (Mbp) | **96.7** | 90.2 | 69.7 |
| NGA50 (Mbp) | **96.6** | 75.1 | 69.7 |
| NGA75 (Mbp) | **44.0** | 36.4 | 30.5 |

